# Seven new Neocallimastigomycota genera from fecal samples of wild, zoo-housed, and domesticated herbivores: Description of *Ghazallomyces constrictus* gen. nov., sp. nov., *Aklioshbomyces papillarum* gen. nov., sp. nov., *Agriosomyces longus* gen. nov., sp. nov., *Capellomyces foraminis* gen. nov., sp. nov. and *Capellomyces elongatus* sp. nov., *Joblinomyces apicalis* gen. nov., sp. nov., *Khoyollomyces ramosus* gen. nov., sp. nov., and *Tahromyces munnarensis* gen. nov., sp. nov.

**DOI:** 10.1101/642694

**Authors:** Radwa A. Hanafy, Vikram B. Lanjekar, Prashant K. Dhakephalkar, Tony M. Callaghan, Sumit S. Dagar, Gareth W. Griffith, Mostafa S. Elshahed, Noha H. Youssef

## Abstract

We isolated and characterized sixty-five anaerobic gut fungi (AGF, Neocallimastigomycota) strains from fecal samples of five wild (W), one zoo-housed (Z), and three domesticated (D) herbivores in the US states of Texas (TX) and Oklahoma (OK), Wales (WA), and the Indian states of Kerala (KE) and Haryana (HA). Phylogenetic assessment based on D1-D2 region of the large rRNA subunit (LSU) identified seven distinct lineages, with strains recovered from Axis Deer (W-TX) clustering within the *Orpinomyces-Neocallimastix-Pecoramyces-Feramyces* clade; Boer Goat-domesticated Goat strains (W-TX, D-KE) clustering within the *Oontomyces-Anaeromyces-Liebetanzomyces* clade; and domesticated Goat and Sheep strains (D-HA) as well as Nilgiri Tahr strains (W-KE) forming two distinct clades associated with genus *Buwchfawromyces*. The remaining three lineages, represented by strains recovered from Mouflon-Boer Goat (W-TX), White Tailed Deer (W-OK), and Zebra-Horse (Z-OK, and D-WA), displayed no specific suprageneric affiliation. All strains displayed monocentric thalli and produced mono/uniflagellate zoospores with the exception of Axis Deer strains, which produced polyflagellate zoospores. Isolates displayed multiple interesting microscopic features including sporangia with tightly constricted necks and fine septa at the base (Axis Deer), papillated and pseudo-intercalary sporangia (White-Tailed Deer), swollen sporangiophores and zoospores with long flagella (Mouflon-Boer Goat), zoospore release through an apical pore followed by either sporangial wall collapse (Axis Deer and Boer Goat-domesticated Goat) or sporangial wall remaining intact after discharge (Zebra-Horse), multi-sporangiated thalli with branched sporangiophores (Zebra-Horse), and short sporangiophores with subsporangial swellings (Nilgiri Tahr). Internal transcribed spacer-1 region (ITS-1) sequence analysis indicated that Zebra-Horse strains are representatives of the AL1 lineage, frequently encountered in culture-independent surveys of the alimentary tract and fecal samples from hindgut fermenters. The other six lineages, five of which were isolated from wild herbivores, have not been previously encountered in such surveys. Our results significantly expand the genus level diversity within the Neocallimastigomycota, and strongly suggest that wild herbivores represent a yet-untapped reservoir of AGF diversity. We propose the creation of seven novel genera and eight novel Neocallimastigomycota species to accommodate these strains, for which we propose the names *Agriosomyces longus* (Mouflon and wild Boer Goat), *Aklioshbomyces papillarum* (White tailed Deer), *Capellomyces foraminis* (wild Boar Goat) and *C. elongatus* (domesticated Goat), *Ghazallomyces constrictus* (Axis Deer), *Joblinomyces apicalis* (domesticated Goat and Sheep), *Khoyollomyces ramosus* (Zebra-Horse), and *Tahromyces munnarensis* (Nilgiri Tahr). The type species are strains Axs-31, WT-2, MS-4, BGB-11, GFKJa1916, GFH683, ZS-33, and TDFKJa193, respectively.

## INTRODUCTION

Members of the anaerobic gut fungi (AGF, Phylum Neocallimastigomycota) colonize the alimentary tract of mammalian and reptilian herbivores (Gruninger and others 2014; Ljungdahl 2008). Interest in the taxonomy, ecology, cell biology, and genomics of the Neocallimastigomycota has been driven by their unique habitat, physiological preferences, and evolutionary history, as well as by their possession of superior plant polymers degradation capacities (Youssef and others 2013). Such characteristics render them a promising platform for biofuel and biogas production from plant biomass (Ranganathan and others 2017; Young and others 2018).

Currently, eleven different AGF genera have been described (Ariyawansa 2015; Barr and others 1989; Breton and others 1990; Callaghan and others 2015; Dagar and others 2015; Gold and others 1988; Hanafy and others 2017; Hanafy and others 2018; Heath and others 1983; Joshi and others 2018; Li 2016; Ozkose and others 2001). However, it is reasonable to assume that multiple novel, yet-uncultured AGF lineages remain to be isolated and characterized. The inherent difficulty in isolating and maintaining these strictly anaerobic and senescence-prone organisms severely hampers isolation and characterization efforts, and limits the number of research groups dedicated to uncovering AGF diversity. Further, it is entirely plausible that multiple AGF taxa are extremely fastidious, with complex nutritional requirements that are not satisfied in current isolation protocols. Indeed, culture-independent diversity surveys utilizing the internal transcribed spacer 1 (ITS-1) as a phylogenetic marker demonstrated that multiple novel AGF lineages remain to be isolated and characterized (Kittelmann and others 2012; Liggenstoffer and others 2010; Paul and others 2018; Mura and others 2019).

On the other hand, culturing efforts have occasionally recovered novel AGF strains that bear no clear similarities to clades identified in culture-independent studies (Callaghan and others 2015; Joshi and others 2018). This surprising observation could be attributed to mismatches in the isolates’ ITS-1 region to commonly utilized ITS-1 AGF primers (Callaghan and others 2015), extremely narrow host range of some AGF taxa (Callaghan and others 2015), or existence in extremely low relative abundance *in-situ*.

Moreover, it is important to note that while both culture-based and culture-independent surveys have reported the presence of AGF communities in a relatively wide range of animal hosts, such studies by no means represent an exhaustive catalogue of global AGF diversity in nature. For example, due to logistical considerations, the great majority of studies have utilized samples from domesticated herbivores, with efforts to isolate AGF strains from wild herbivores being extremely rare (Nagpal and others 2011; Paul and others 2010; Tuckwell and others 2005; Hanafy and others 2018).

In an effort to broaden the current Neocallimastigomycota global culture collection, we conducted a multi-year isolation effort targeting novel AGF taxa in fecal samples from a wide range of wild, zoo-housed, and domesticated herbivorous mammals. Here, we report on the isolation and characterization of seven novel AGF genera, including the first cultured representative of the hitherto uncultured AGF lineage AL1. The results expand the known AGF genus-level diversity by >50% (from 11–18), and strongly suggest that wild undomesticated herbivores represent a yet-untapped reservoir of novel AGF taxa.

## MATERIALS AND METHODS

### Samples

Fecal samples were obtained from Axis Deer (*Axis axis*), White Tailed Deer (*Odocoileus virginianus*), Mouflon Sheep (*Ovis orientalis*), and Boer Goat (*Capra aegagrus*) in two separate hunting expeditions in Sutton and Val Verde counties (TX), and Payne county (OK) in October 2017 and April 2018 (TABLE 1). The hunting parties had all appropriate licenses, and the animals were shot either on private land with the owner’s consent or on public land during the hunting season. Samples were also obtained from a Grevy’s Zebra (*Equus grevyi*) housed in the Oklahoma City Zoo in May 2018, with the sampling protocol approved by the Oklahoma City Zoo and Botanical Garden’s Scientific Review Committee. All fecal samples were placed on ice on site, transferred to the laboratory within 24h of collection, where they were immediately used as an inoculum for subsequent enrichment and isolation procedures.

In India, dried fecal samples were obtained from Nilgiri Tahr (*Nilgiritragus hylocrius*), and domesticated but forest grazing Goat (*Capra aegagrus hircus*) in Munnar in the State of Kerala. Fresh fecal samples were also obtained from domesticated Goats and Sheep (*Ovis aries*) in Sonipat in the State of Haryana. Fresh fecal samples were transferred to the laboratory within 24h of collection, while dried fecal samples were transferred within 72 hours of collection. In Wales, samples were obtained from two domesticated Horses in Llanbadarn Fawr, Ceredigion County, and promptly transferred to the laboratory for processing.

### Isolation procedures

In the USA, isolation of anaerobic fungal strains was conducted as previously described (Hanafy and others 2017). Feces were suspended in rumen-fluid (RF) media (Calkins and others 2016; Hanafy and others 2018) with either cellobiose, or 0.5% cellobiose and switchgrass (0.5%) used as a substrate. Antibiotics (50 μg/mL kanamycin, 50 μg/mL penicillin, 20 μg/mL streptomycin, and 50 μg/mL chloramphenicol, respectively) were added to inhibit growth of bacteria. Samples were serially diluted and incubated at 39 C for 24–48 h. Dilutions showing visible signs of growth (clumping or floating plant materials and production of gas bubbles) were then used for the preparation of roll tubes (Hungate 1969) on RF-cellobiose agar media. Single colonies were picked into liquid RF-cellobiose media, and at least three rounds of tube rolling and colony picking were conducted to ensure purity of the obtained colonies. Strains were maintained by bi-weekly subculturing into RF-cellobiose media. In India, the fecal samples were homogenised in anaerobic diluent (McSweeney and others 2005) using BagMixer (Interscience, France). One ml of homogenate was inoculated into 9 ml fungal culture medium (Joshi and others 2018) containing neutral detergent fibre (NDF) as the sole carbon source, and serially diluted up to 10^−3^ dilution. The antibiotics benzylpenicillin and streptomycin sulfate (final concentration 2 mg/ml) were used to inhibit the bacterial growth. Following incubation at 39±1 C for 5–10 d, the tubes showing visible colonization of NDF were used to isolate pure cultures of anaerobic fungi using serum roll bottle method as described previously (Joshi and others 2018). The colonies differing in morphology were picked, grown in liquid culture medium, and re-roll bottled until single culture was established.

Long-term storage was conducted by surface inoculation of RF-cellobiose agar media as described previously (Calkins and others 2016), or by cryopreservation at −80 °C using 0.64 M ethylene glycol as the cryoprotectant (Callaghan and others 2015). Cultures are available at Oklahoma State University, Department of Microbiology and Molecular Genetics culture collection, and at MACS Collection of Microorganisms (MCM), Agharkar Research Institute, Pune, India. In Wales, isolation procedures were conducted as previously described in (Callaghan and others 2015).

### Morphological characterization

The colony morphology of 3d old cultures on roll bottles was measured using a stereomicroscope (Leica M205 FA) equipped with a digital camera (Leica DFC450 C), or directly from the roll tube. Samples for light and scanning electron microscopy were obtained from liquid cultures at various stages of growth. Lactophenol cotton blue stain was used for visualization of fungal structures using an Olympus BX51 microscope (Olympus, Center Valley, Pennsylvania) equipped with a DP71 digital camera (Olympus), or a phase contrast microscope equipped with a Canon DS126191 digital camera. For examination of nuclei localization, samples were stained with 4′, 6 diamidino-2-phenylindole (DAPI, 10 μg/ml), as previously described (Callaghan and others 2015; Hanafy and others 2018; Joshi and others 2018) and examined using a fluorescence Olympus BX51 microscope (Olympus, Center Valley, Pennsylvania) equipped with a Brightline DAPI high-contrast filter set for DAPI fluorescence and a DP71 digital camera (Olympus), or an Olympus BX53 differential interference contrast (DIC) microscope equipped with a DP73 digital camera (Olympus). Scanning electron microscopy was conducted with a FEI quanta scanning electron microscope (Hillsboro, Oregon, USA), or a Carl Zeiss EVO MA15 (Hanafy and others 2017; Joshi and others 2018).

### Phylogenetic analysis

Biomass was harvested and crushed in liquid N_2_. DNA was extracted from the ground fungal biomass using DNeasy PowerPlant Pro Kit (Qiagen Corp., Germantown, MD, USA) according to the manufacturer’s instructions, or using the CTAB DNA extraction protocol (Joshi and others 2018). To assess phylogenetic relationships, the ITS-1 region, and the D1/D2 region of the 28S rRNA (hereafter LSU) were amplified using the MN100 (5′-TCCTACCCTTTGTGAATTTG-3′) / MNGM2 (5′-CTGCGTTCTTCATCGTTGCG-3′) pair, and the NL1 (5′-GCATATCAATAAGCGGAGGAAAAG-3′) / NL4 (5′-GGTCCGTGTTTCAAGACG G-3′) pair, respectively as previously described (Hanafy and others 2017; Joshi and others 2018). The resulting PCR amplicon for the ITS-1 region was cloned into a TOPO-TA cloning vector according to the manufacturer’s instructions (Life Technologies®, Carlsbad, CA) and several clones were Sanger sequenced, while the purified LSU PCR amplicons were directly sequenced using the services of the Oklahoma State University DNA core facility or a commercial provider (1^st^ BASE, Singapore). The obtained sequences were aligned to anaerobic fungal reference ITS-1 and LSU sequences downloaded from GenBank using MAFFT aligner (Nakamura and others 2018) and the alignments were used to construct maximum likelihood phylogenetic trees in MEGA7 (Kumar and others 2016), using *Chytriomyces* sp. JEL176 as the outgroup. Bootstrap values were calculated on the basis of 100 replicates.

### Ecological distribution

We queried GenBank and ITS-1 datasets (Kittelmann and others 2012; Liggenstoffer and others 2010; Paul and others 2018) using reference ITS-1 sequences from strains recovered in this study. The phylogenetic position of all closely related sequences (> 87% sequence similarity) was evaluated by insertion into maximum likelihood trees. Taxonomy of uncultured taxa followed the schemes outlined in prior publications (Kittelmann and others 2012; Liggenstoffer and others 2010; Paul and others 2018).

### Accession numbers

Sequences generated in this study have been deposited in GenBank under accession numbers MK881965-MK882046, MK775304, MK775310-MK775313, MK775315, MK775321-MK775324, MK755326-MK755327, MK755330, MK755333. Alignments and phylogenetic trees are available through TreeBase under study accession URL http://purl.org/phylo/treebase/phylows/study/TB2:S24394

## RESULTS

### Isolation summary

Sixty-five different strains were obtained and characterized in this study (TABLE 1). These isolates were obtained from fecal samples of five wild undomesticated herbivores: Axis Deer (W-TX), White tailed Deer (W-OK), Mouflon (W-TX), Boer Goat (W-TX), and Nilgiri Tahr (Munnar, Kerala, India), one zoo-housed Grevy’s Zebra (Z-OK), two domesticated Horses (D-WA), two domesticated Goats (D-HA and D-KE) and a domesticated Sheep (D-HA) (TABLE 1). Morphological, microscopic and phylogenetic analysis described below grouped these isolates into seven distinct clades (Labeled clades 1-7 in TABLE 1 according to alphabetical order of suggested genus names). Isolates representing three clades were obtained from one host animal only: White Tailed Deer strains (Clade 2), Axis Deer strains (Clade 4), and Nilgiri Tahr strains (clade 7); while isolates representing four clades were identified in more than sample: Mouflon-Boer Goat strains (clade 1), Boer Goat-domesticated Goat strains (clade 3), domesticated Sheep-Goat strains (clade 5), and Zebra-Horse strains (clade 6). No specific morphological or microscopic decipherable differences were identified between different strains belonging to most of these clades, and one strain from each group was chosen for detailed analysis (TABLE 1). The only two exceptions were: 1. Strains belonging to clade 3 (Boer Goat-domesticated Goat); where strains from wild Boer Goat (W-TX) displayed distinct microscopic and phylogenetic differences from those obtained from the domesticated Goat (D-KE) to warrant the detailed characterization and eventual description of two different strains (TABLE 1 and detailed descriptions below), and 2. Strains belonging to clade 6 (Zebra-Horse) where few microscopic, but negligible phylogenetic differences were observed between the 16 Zebra strains (Z-OK) and the 5 Horse strains (D-WA) identified. These differences are highlighted below, but we do not believe they warrant the description of a new species, given the negligible sequence divergences between these strains. Below, we provide a detailed characterization of these seven novel groups.

### Colony morphology and macroscopic growth characteristics

Clade 1 (Mouflon-Boer Goat) strain MS-2 produced small brown circular colonies (0.2–1 mm diam.) on agar, and a thin biofilm-like growth in liquid media (FIG. 1a). Clade 2 (White tailed Deer) strain WT-2 produced beige circular colonies (0.5–2.5 mm diam.) with a brown central core of dense sporangial structures and an outer ring of light gray hyphal growth. In liquid media, it produced heavy growth of thick biofilms that firmly attached to the tube’s glass surface (FIG. 1b). Clade 3 (Boar Goat-domesticated Goat) strain BGB-11 produced small circular brown colonies (0.1–0.5 mm diam.), with dark center of sporangia structures and a thin fungal biofilm in liquid media (FIG. 1c), while Boar Goat-domesticated Goat strain GFKJa1916 produced compact cottony off-white colonies of 2–3 mm size, with a fluffy center of thick sporangial structures and surrounded by radiating rhizoids (FIG. 1d). In liquid media, strain GFKJa1916 produced numerous fungal thalli attached to the glass bottles on initial days of growth, which later developed into thin mat-like structures. Clade 4 (Axis Deer) strain Axs-31 produced small circular white colonies (1–4 mm diameter), with a brown central core of dense sporangial structures on agar and a thick fungal biofilm-like growth in liquid media (FIG. 1e). Clade 5 (Domesticated Goat-Sheep) strain GFH683 produced 1–2 mm sized colonies, having a dense dark central core of abundant sporangial growth, surrounded by long and thin radiating rhizoids (FIG. 1g). In liquid media, strain GFH683 produced numerous fungal thalli attached to the glass bottles on initial days of growth, which later developed into thin mat-like structures. Clade 6 (Zebra-Horse) strains Zebra strain ZS-33 produced small yellow to yellowish brown irregularly shaped colonies (FIG. 1f). In liquid media, the fungal thalli were loose and exhibited a sand-like appearance resembling liquid growth patterns generally observed with isolates belonging to the bulbous genera *Caecomyces* and *Cyllamyces* (personal observation) (FIG. 1f). Finally, clade 7 (Nilgiri Tahr) strain TDFKJa193 were smaller, approximately 1 mm in size, white in color with a compact and fluffy center, surrounded by dotted circles of fungal thalli. In liquid media, the strain produced numerous fungal thalli attached to the glass bottles on initial days of growth, which later developed into thin mat-like structures (FIG. 1h).

**FIG. 1.**
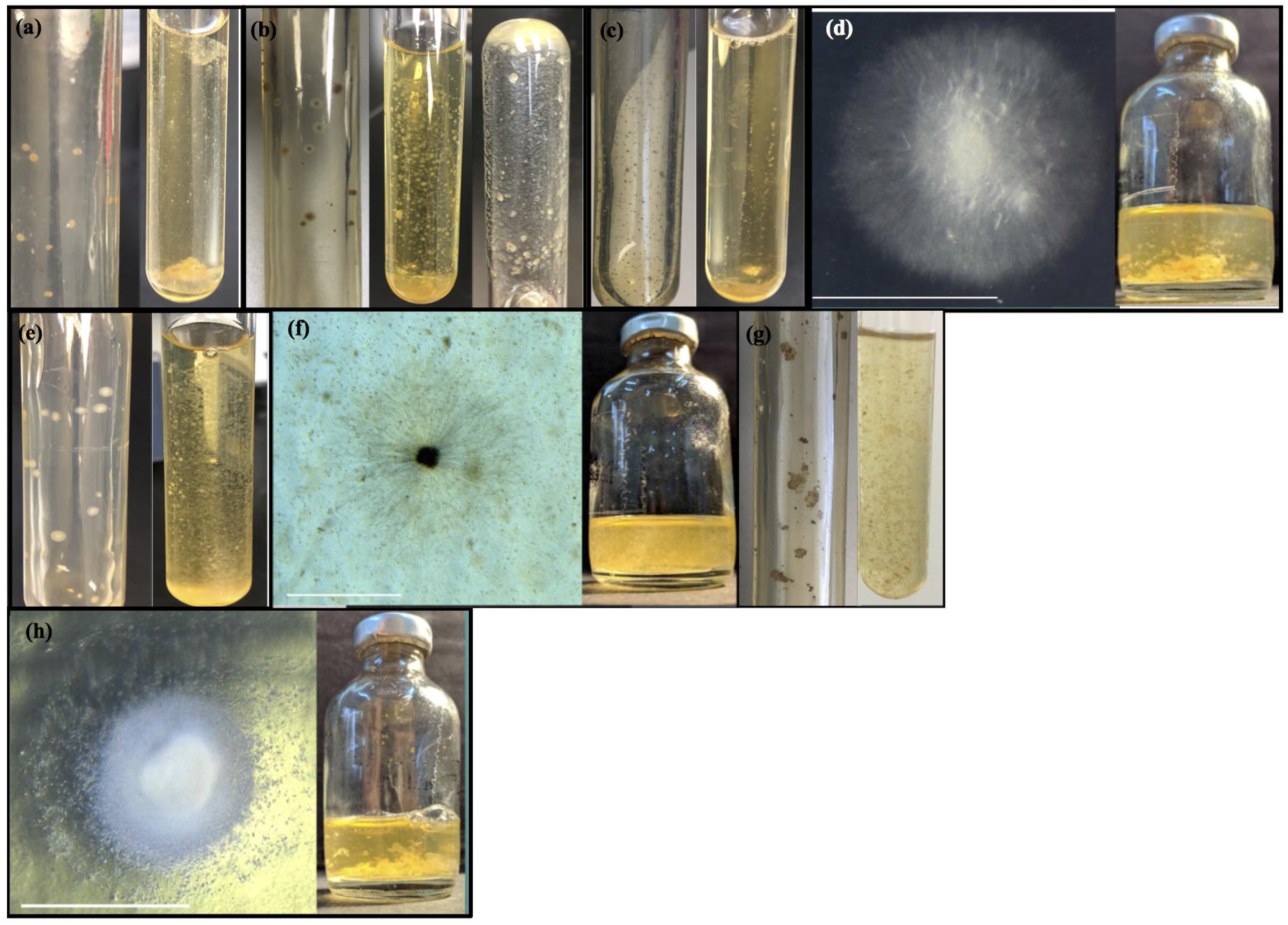
Macroscopic features of colony morphology on agar media and fungal biomass in liquid media for (a) *Agriosomyces longus* strain MS2, brown circular colonies and thin fungal biofilm in liquid medium. (b) *Aklioshbomyces papillarum* strain WT-2, beige circular colonies with a brown central core of sporangia and thick fungal biofilm in liquid medium that firmly attaches to the tube’s glass surface, note the inverted tube shows how the fungus thalli attached to the tube. (c) *Capellomyces foraminis* strain BGB-11, small brown circular colonies with dark center and thin fungal biofilm in liquid medium. (d) *Capellomyces elongatus* strain GFKJa1916, compact cottony off-white circular colonies with a fluffy central core of sporangia surrounded by radiating sporangia and thin fungal biofilm in liquid medium that attach to the tube’s glass. (e) *Ghazallomyces constrictus* strain Axs-31, white circular colonies on roll tubes and thick fungal biofilm in liquid medium. (f) *Joblinomyces apicalis* strain GFH683, beige circular colonies with a brown central core of sporangia and thin fungal biofilm in liquid medium. (g) *Khyollomyces ramosus* strain ZS-33, yellowish brown colonies of irregular shape on agar medium and loose fungal thalli with sand-like appearance. (h) *Tahromyces munnarensis* strain TDFKJa193 white circular colonies with fluffy center, surrounded by dotted circles of fungal thalli and thick fungal biofilm in liquid medium that attach to the tube’s glass.

### Microscopic features

#### Clade 1 (Mouflon-Boer Goat) strains

Strain MS2 produced small globose zoospores, with an average diameter of 4 ± 1.1 μm (average ± standard deviation values from 29 zoospores, range 2.7–7.5 μm). Zoospores were mainly mono-flagellate with a flagellum length of 22 ± 3.8 μm (average ± standard deviation values for 29 zoospores, range 16.6–30 μm), approximately 5–6 times longer than the zoospore body (FIG. 2a). Biflagellate zoospores (FIG. 2b) were rarely encountered. Zoospores germinated into monocentric thalli with filamentous anucleate rhizoidal systems (FIG. 2c-d). Both endogenous and exogenous sporangia were observed, which were very homogenous and displayed no pleomorphism. Endogenous sporangia were globose, with a diameter range of 15–65 μm (FIG. 2e-f). The rhizoid was swollen below the sporangial neck, which was tightly constricted (FIG. 2e-f). Exogenous sporangia were also consistently globose and developed at the end of swollen sporangiophores (30–80 μm L X 5–10 μm W) (FIG. 2g-h). The sporangial neck was constricted with a narrow neck port. Zoospores were released through dissolution and rupturing of the sporangial wall (FIG. 2i).

**FIG. 2.**
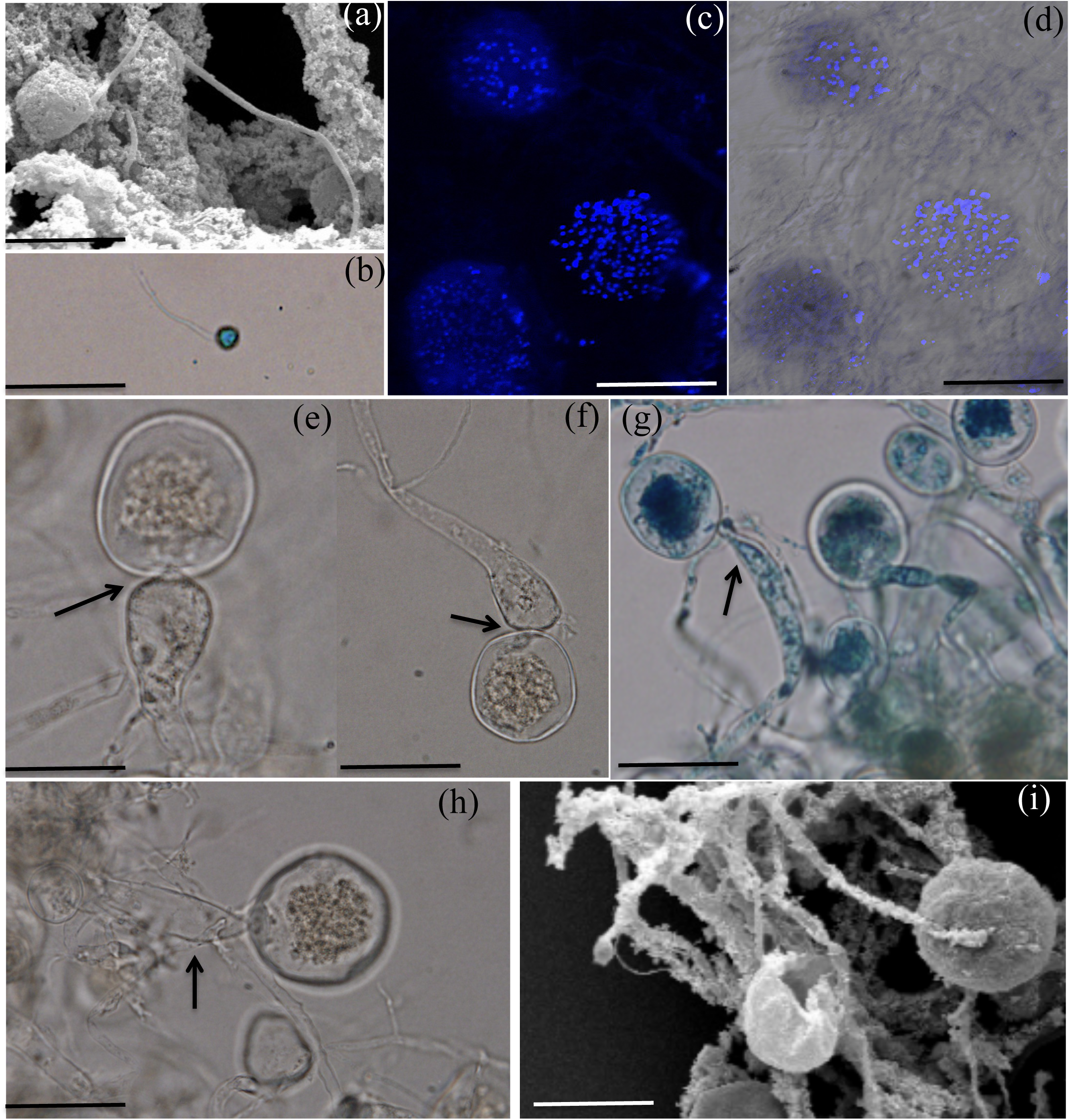
Microscopic features of *Agriosomyces longus* (Clade 1, Mouflon-Boer Goat) strain MS2. Light (e-h) and scanning electron (a and i) micrographs are shown. DAPI staining for nuclei visualizing using a fluorescence microscope equipped with a Brightline DAPI high-contrast filter set (c). Overlay image is shown in (d). (a) A monoflagellate zoospore. (b) A biflagellate zoospore. (c-d) Monocentric thalli, with nuclei occurring in sporangia, not in rhizoids or sporangiophores. (e-f) Endogenous globose sporangia with tightly constricted necks and subsporangial swellings (arrows). (g-h) Exogenous globose sporangia, note the swollen sporangiophores. (i) An empty sporangium after zoospore release and rupturing of the sporangial wall. Bar: a=5 μm, b=20 μm c-i=50 μm.

#### Clade 2 (White-tailed Deer) strains

Strain WT-2 produced globose zoospores, with an average diameter of 7.4 ± 2.4 μm (average ± standard deviation values for 35 zoospores, range 4.5–13 μm). Zoospores were mostly monoflagellate, with an average flagellum length of 22.8 ± 6.3 μm (average ± standard deviation values for 35 zoospores, range 12–35 μm) (FIG. 3a). Zoospores with two (FIG. 3b) to three (FIG. 3c) flagella were less frequently observed. Fungal thalli were consistently monocentric with filamentous anucleate rhizoids (FIG. 3d). Germination of zoospores produced two types of monocentric thalli, endogenous and exogenous. Endogenous sporangia with single (FIG. 3e) and two adjacent rhizoidal systems (FIG. 3f) were observed. Occasionally, pseudo-intercalary endogenous sporangia (sporangia present in the middle of two main rhizoids) were encountered (FIG. 3g), Similar to what have previously been observed with the genera *Oontomyces* (Dagar and other, 2015) and *Feramyces* (Hanafy and others, 2017). Exogenous sporangia developed at the end of unbranched sporangiophores of varying length from a few microns to 230 μm (FIG. 3h-j). No morphological differences were noticed between endogenous and exogenous sporangia and their shapes ranged from ovoid (FIG. 3e-f), globose (FIG. 3g-h), obpyriform (FIG. 3j-k and 3n-o), and ellipsoidal (FIG. 3i and 3l). Many, but not all, sporangia were papillated with one (FIG. 3m-p) or two (FIG. 3q) papillae. These papillated sporangia are similar to those previously observed in *Piromyces mae* (Li and others 1990). It is believed that these papillae disintegrate to facilitate zoospore release. However, we were unable to observe zoospore discharge through papillae in strain WT-2.

**FIG. 3.**
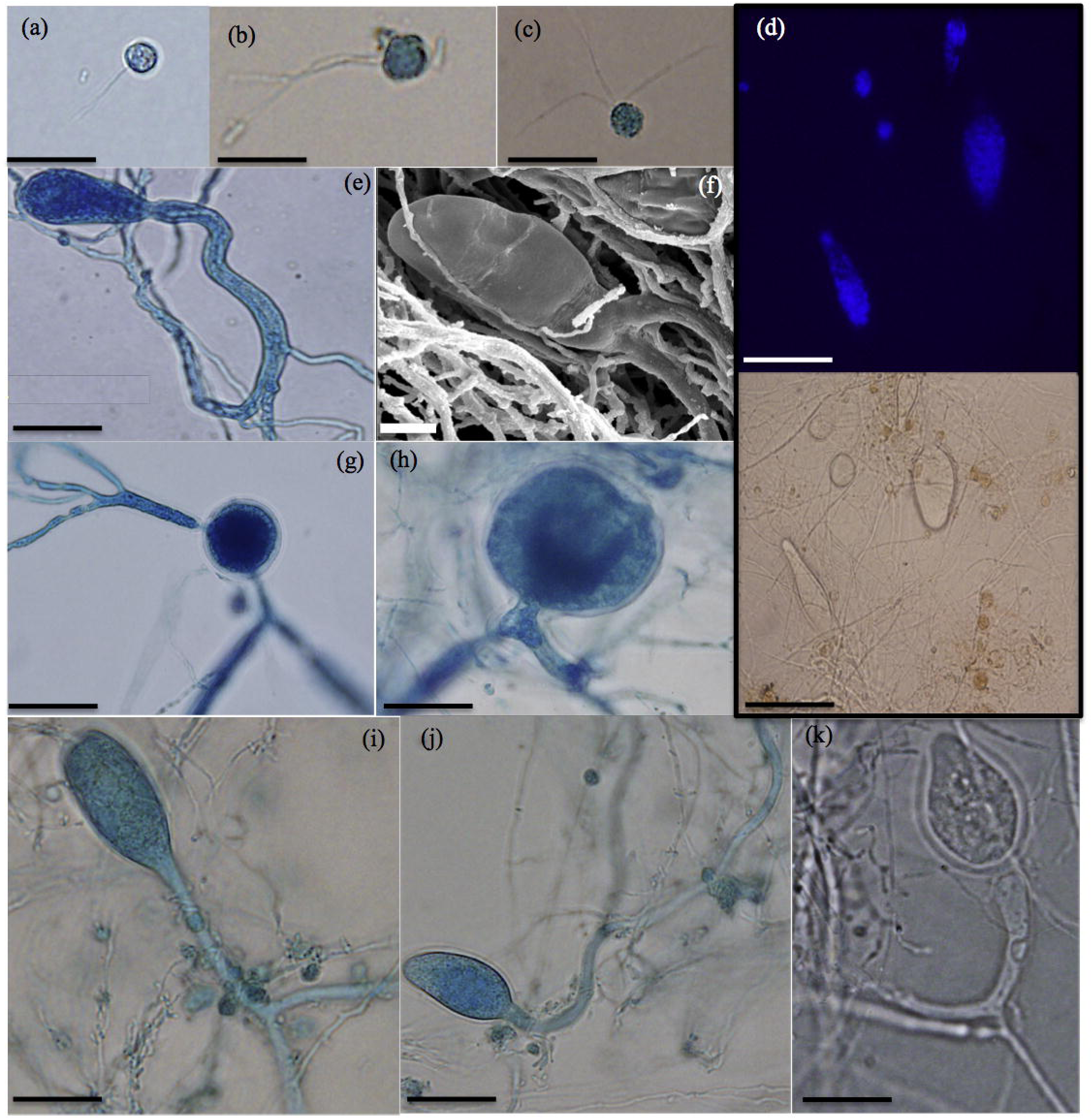
Microscopic features of *Aklioshbomyces papillarum* (Clade 2, White-Tailed Deer) strain WT-2. Light (a-e, and g-q) and scanning electron (f) micrographs are shown. Light microscopy pictures were examined after staining with lactophenol cotton blue (a-c, e, and g-q), as well as following nuclei staining with DAPI (d). (a) A monoflagellate zoospore. (b) A biflagellate zoospore. (c) A triflagellate zoospore. (d) Monocentric thalli, with nuclei occurring in sporangia, not in rhizoids or sporangiophores. (e-g) Endogenous sporangial development: (e) Ovoid sporangium with single rhizoidal system, (f) Ovoid sporangium with two main rhizoidal systems, (g) Globose pseudo-intercalary sporangium, between two main rhizoidal systems. (h-k) Exogenous sporangial development: (h) Globose sporangium on a very short sporangiophore, (i) Ellipsoidal sporangium, (j) Obpyriform sporangium on a long sporangiophore, (k) Obpyriform sporangium. (l) Ellipsoidal sporangium. (m) Sporangia with lateral single papilla, (n-p) Sporangia with terminal single papilla. (q) Sporangium with two papillae. Bar: a-c, e, f, n & o =20 μm, g-k, p & q =50 μm, d, l & m =100 μm.

#### Clade 3 (Boer Goat-domesticated Goat) strains

Boer Goat Strain BGB-11 produced globose zoospores, with an average diameter of 5.5 ± 0.97 μm (average ± standard deviation values for 40 zoospores, range 4–7 μm). The majority of zoospores were mono-flagellate with a flagellum length of 19.6 ± 3.2 μm (average ± standard deviation values for 40 zoospores, range 15–25 μm), (FIG. 4a). Occasionally, biflagellate zoospores were observed (FIG. 4b). Zoospores encystment followed flagellar shedding (FIG. 4c). Zoospore cyst germinated producing germ tube (FIG. 4d) that subsequently branched (FIG. 4e) into monocentric thalli with filamentous anucleate rhizoidal systems (FIG. 4f-g).

**FIG. 4.**
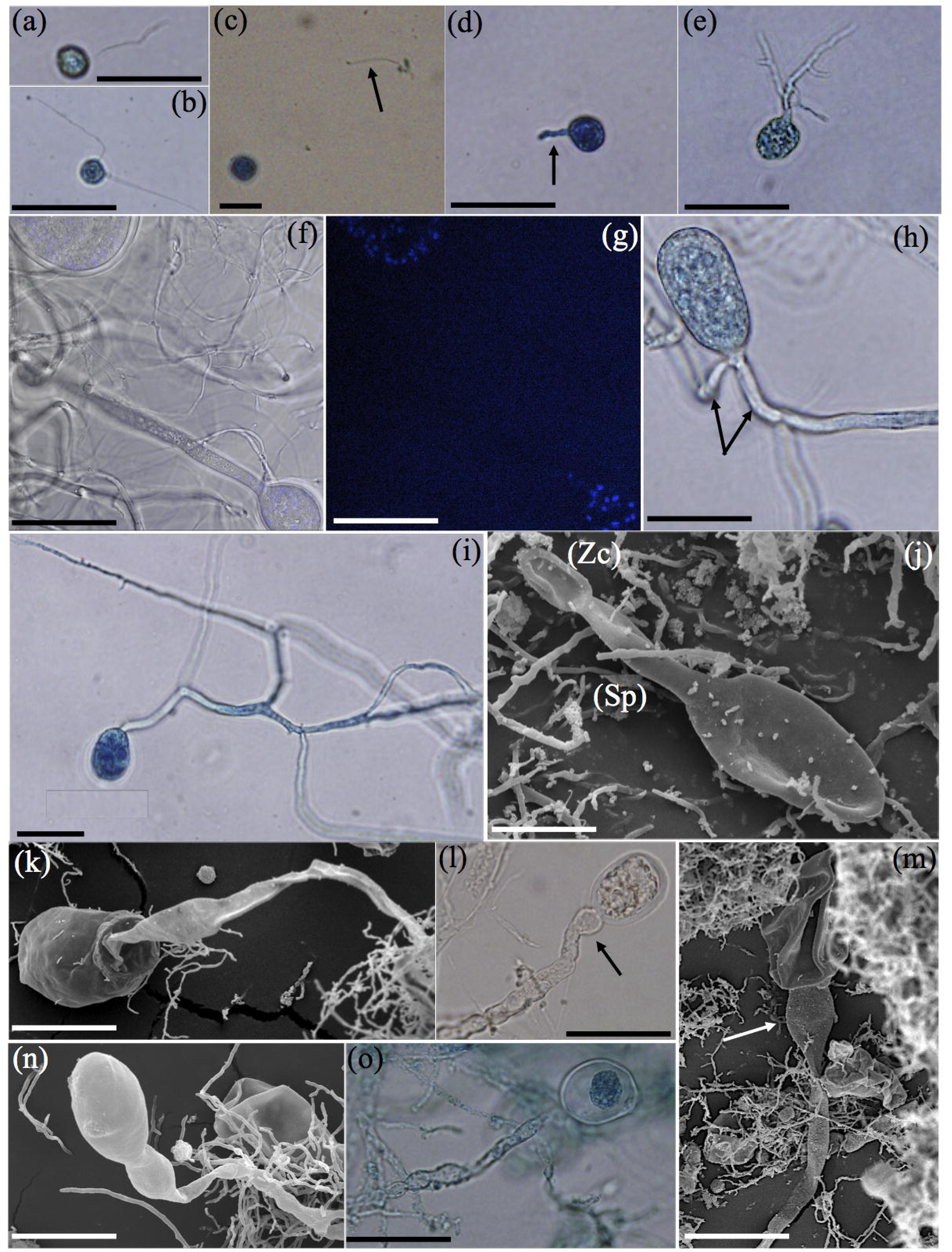
Microscopic features of *Capellomyces foraminis* (Clade 3, Boer Goat) strain BGB-11. Light (a-i, l, o & p) and scanning electron (j, k, m, n, q &r) micrographs are shown. Light microscopy pictures were examined after staining with lactophenol cotton blue (a-e, h, i, & o), as well as following nuclei staining with DAPI (f). Overlay image is shown in (g). (a) A monoflagellate zoospore. (b) A biflagellate zoospore. (c) Zoospore cyst, arrow points to the shed flagellum. (d) Germinating zoospore cyst producing a germ tube (arrow). (e) Rhizoidal system development. (f-g) Monocentric thalli with nuclei occurring in sporangia, not in rhizoids or sporangiophores. (h-i) Endogenous sporangial development: (h) Ellipsoidal sporangium with two main rhizoidal systems (arrows), (i) Ovoid sporangium with single rhizoidal system. (j-q) Exogenous sporangial development: (j) Exogeous sporangium with a short sporangiophore (Sp), note the empty zoospore cyst (Zc). (k) Ovoid sporangium with a long sporangiophore, (l) Ovoid sporangium on a long sporangiophore ends with sub-sporangial swelling (arrow), (m) Collapsed empty sporangium on a long sporangiophore ends with sub-sporangial swelling (arrow), (n) Constricted ellipsoidal sporangium, (o-p) Globose Sporangia. (q) Zoospores ae released through apical pore. (r) An empty sporangium following zoospores release. Abbreviations: (Sp), sporangiophore; (Zc), zoospore cyst. Bar: a-j, l, o & p =20 μm, k, m, n, q & r =50 μm.

The expansion of the zoospore cysts resulted in the formation of endogenous sporangia that were ellipsoidal (FIG. 4h) and ovoid (FIG. 4i). In addition to endogenous sporangia, exogenous sporangia were also observed at the end of unbranched sporangiophores ranging in length between 20–150 μm (FIG. 4j-p). Some of the sporangiophores ended with sub-sporangial swellings (FIG. 4l-m). Exogenous sporangia varied in shape from ovoid (FIG. 4k-l), ellipsoidal with a single constriction (FIG. 4n), and globose (FIG. 4o-p). Zoospores were liberated through a wide apical pore at the top of the sporangia followed by sporangial wall collapse (FIG. 4m, q-r).

Domesticated Goat strain GFKJa1916 on the other hand produced globose zoospores (FIG. 5a), with an average diameter of 4–5 μm. The majority of zoospores were mono-flagellate with a flagellum length of 15–20 μm. Bi- and tri-flagellate zoospores were also observed. Strain GFKJa1916 zoospores germinated either endogenously or exogenously into a single monocentric thallus, which was also confirmed by the presence of nuclei only in sporangia and their absence in rhizoids (FIG. 5b-c). Endogenous sporangia varied in shape between cylindrical, elongate, globose, sub-globose, ellipsoid & obovoid with sizes ranging between 8–60 μm wide & 10–140 μm long (FIG. 5d-g). Unlike Boer Goat Strain BGB-11, exogenous sporangia in the domesticated Goat strain GFKJa1916 developed at the end of long thick sporangiophores (up to 300 µm in some cases) (FIG. 5h-l), and multisporangiate thalli were commonly observed with two sporangia of either the same (FIG. 5j) or different (FIG. 5K-l) shape, similar to *Piromyces rhizinflatus* (Ho and Barr 1995) and *Neocallimastix frontalis* (Barr and others 1995).

**Fig 5.**
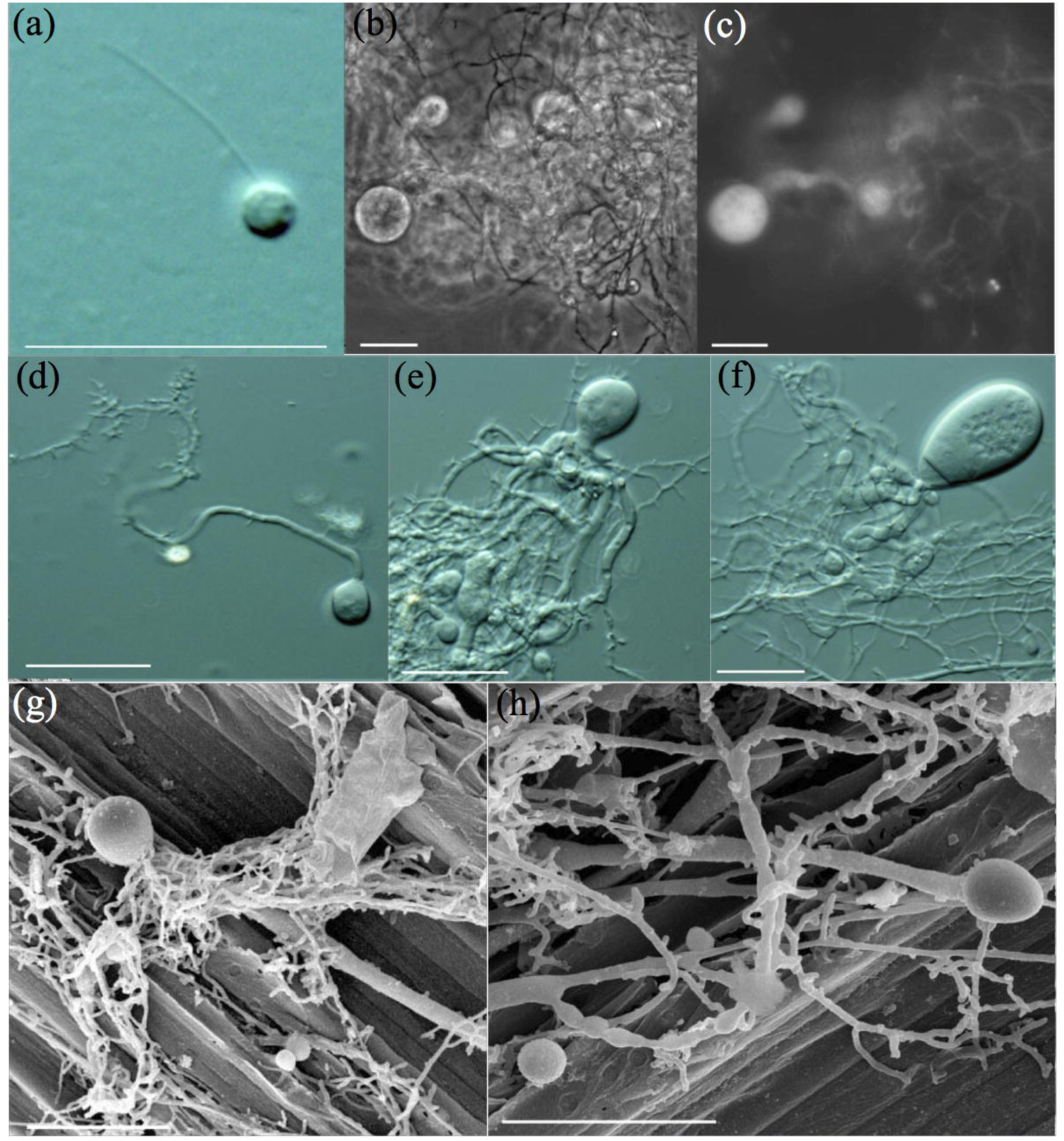
Microscopic features of *Capellomyces elongatus* (Clade 3, domesticated Goat) strain GFKJa1916. Differential interference contrast (a, d-f and i-l), scanning (g-h), phase contrast (b) and fluorescence (c) micrographs. (a) A monoflagellate zoospores. (b-c) Monocentric thalli; nuclei were observed in sporangia, not in rhizoids or sporangiophores. (d-g) Endogenous sporangia. (d) Globose endogenous sporangium with one main rhizoidal systems. (e-f) Endogenous sporangia with multiple rhizoidal systems. (g) Endogenous sporangium on wheat straw fibers. (h-l) Exogenous sporangia: (h) ovoid-shaped sporangium with long sporangiophore. (i) Multiple ovoid, and globose exogenous sporangia with long sporangiophores. (j) Multisporangiate thallus with two sporangia (same shape). (k-l) Multisporangiate thallus with two sporangia (different shape). Scale bar = 20 µM.

#### Clade 4 (Axis Deer) strains

Strain Axs-31 produced globose zoospores, with an average diameter of 8.1 ± 1.3 μm (average ± standard deviation values for 35 zoospores, range 6–10.5 μm). All zoospores were polyflagellate, with (7–14) flagella and an average flagellum length of 23.5 ± 4.9 μm (average ± standard deviation values for 35 zoospores, range 16–31 μm) (FIG. 6a). Zoospores germinated into monocentric thalli with highly branched anucleate rhizoidal systems (FIG. 6b-c).

**FIG. 6.**
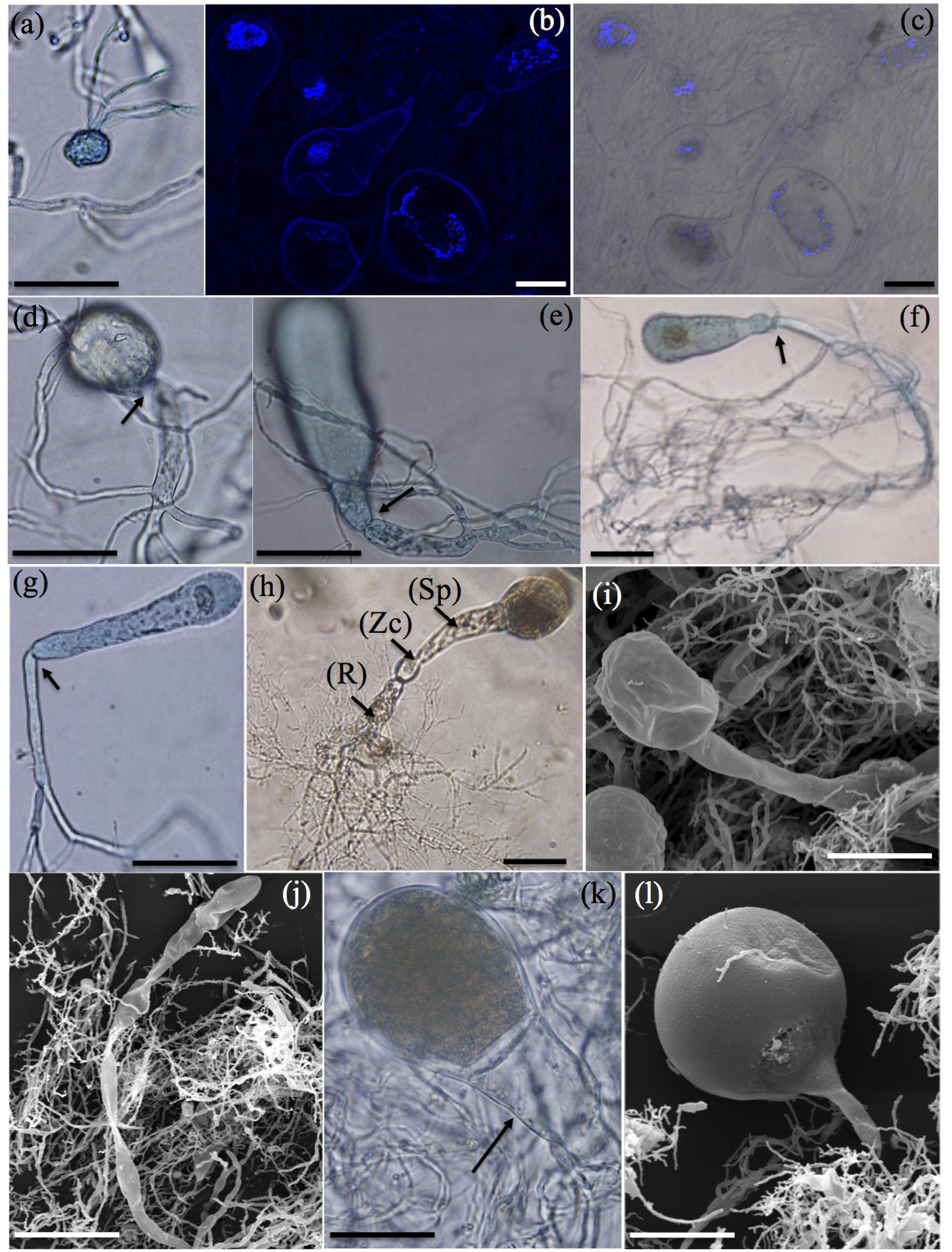
Microscopic features of *Ghazallomyces constrictus* (Clade 4, Axis Deer) strain Axs-31. Light (a-h, f, k and m-p) and scanning electron (i, j, l and q) micrographs are shown. Light microscopy pictures were examined after staining with lactophenol cotton blue (a, d–h, k, and m-p), as well as following nuclei staining with DAPI (b). Overlay image is shown in (c). (a) A polyflagellate zoospore. (b-c) Monocentric thalli, with nuclei occurring in sporangia, not in rhizoids or sporangiophores. (d-g) Endogenous sporangia with tightly constricted necks (arrows): (d) Young globose sporangium, (e) Young tubular sporangium, (f) Mature clavate sporangium, (g) and Mature ellipsoidal sporangium. (h-p) Exogenous sporangia: (h) Young sporangium on a short flattened sporangiophore (Sp), note the persistent empty zoospore cyst (Zc) and the rhizoidal system (R), (i) Ovoid sporangium on short sporangiophore, (j) Ellipsoidal sporangium on long sporangiophore, (k) ovoid sporangium on an eggcup-shaped sporangiophore (arrow), (l) Globose sporangium, (m) Constricted ellipsoidal sporangium with tightly constricted neck (arrow) on long sporangiophore, (n) Pyriform sporangium, note the fine septum at the base of sporangium (arrow), (o) Bowling pin-shaped sporangium, (p) Rhomboidal sporangium with constricted neck and fine septum (arrows), note the persistent empty zoospore cyst (Zc). (q) Zoospores are released through apical pore followed by collapse of the sporangial wall. Abbreviations: (Sp), sporangiophore; (Zc), zoospore cyst; (R), rhizoid. Bar: a=20μm, b-q=50 μm.

Strain Axs-31 exhibited both endogenous and exogenous monocentric thallus development. In endogenous thalli, zoospore cysts enlarged into new sporangia of different shapes, including globose (FIG. 6d), tubular (FIG. 6e), clavate (FIG. 6f) and ellipsoidal (FIG. 6g). Endogenous sporangia displayed tightly constricted necks (point between sporangia and rhizoids) with narrow ports (arrows in FIG. 6d-g).

During exogenous thallus development, zoospore cysts germinated from both ends. Rhizoids developed on one side while sporangiophores developed on the opposite side. The empty zoospore cyst remained as a persistent swollen structure at the base of sporangiophore (FIG. 6h). Exogenous sporangia developed at the end of unbranched sporangiophores of varied lengths. Short sporangiophores had an average length of 6–20 μm (FIG. 6h-i), while long sporangiophores extended up to 200 μm (FIG. 6j). Some of the short sporangiophores had eggcup-shaped appearance (FIG. 6k). Exogenous sporangia were ellipsoidal (FIG. 6j), ovoid (FIG. 6k), globose (FIG. 6l), constricted ellipsoidal (FIG. 6m), pyriform (FIG. 6n), bowling pin-shaped (FIG. 6o), and rhomboidal (FIG. 6p). Sporangial necks were constricted with narrow port (FIG. 6m-p). At maturity, a fine septum developed at the base of the sporangium (FIG. 6n & p, arrow). Zoospores were released through an apical pore followed by collapse of the sporangial wall (FIG. 6q).

#### Clade 5 (Domesticated Goat and Sheep) strains

Strain GFH683 produced globose zoospores (FIG. 8a-b), with an average diameter of 5–6 μm. The majority of zoospores were mono-flagellate with 1, or 2 flagella (FIG. 7a-b). Flagellum length ranged between 20–22 μm. Zoospores germinated to produce both endogenous and exogenous monocentric thalli (FIG. 7c-f), as evident by presence of a single sporangium per thallus, nucleated sporangia but anucleate rhizoids. Endogenous sporangia were globose, sub-globose, ovoid, and obovoid (FIG. 7g) with sizes ranging between 8–40 μm wide & 10–40 μm long. Exogenous sporangia were terminal and varied in shape between globose, ovoid, and obovoid with sporangiophores that varied in length from 20–80 µm (FIG. 7h-i). Zoospores discharge occurred through gradual dissolution of a wide apical portion of sporangial wall, resulting in formation of an empty cup-shaped sporangium (FIG. 7j-l). Such zoospore liberation patterns, and empty cup shaped sporangia were earlier documented for *Piromyces minutus* (Ho and Barr 1995).

**FIG. 7.**
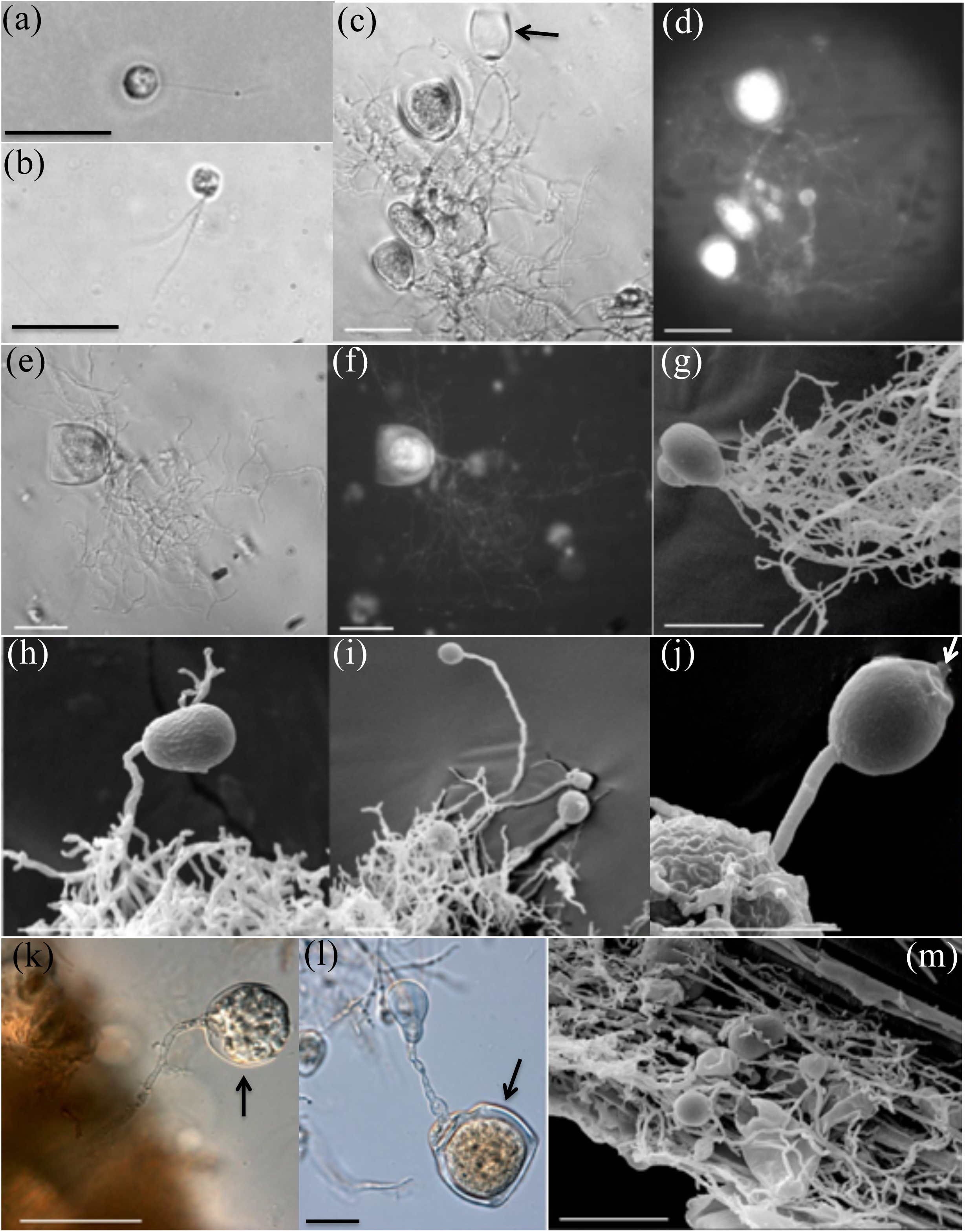
Microscopic features of *Joblinomyces apicalis* (Clade 5, domesticated Goat and Sheep) strain GFH683. Phase contrast (b, c and e), fluorescence (d, f), scanning electron (g-h, and m) differential interference (k-l) micrographs. (a) A monoflagellate zoospore. (b) A biflagellate spherical zoospore. (c-f) Monocentric thalli; nuclei were observed in sporangia, not in rhizoids or sporangiophores, note the empty cup-shaped sporangium after zoospore release (arrow). (g) Ovoid endogenous sporangium. (h-i) Exogenous sporangia: (h) Ovoid sporangium with short sporangiophore. (i) Subglobose sporangium with long sporangiophore. (j-l) Zoospore release: (j) Dissolution of the apical portion of sporangial wall (arrow). (k-l) Cup-shaped sporangia with wide apical pores and intact sporangial walls (arrow). (g) Colonization of rice straw fibers by fungal rhizoids and emanating. Scale bar = 20 µM.

**FIG. 8.**
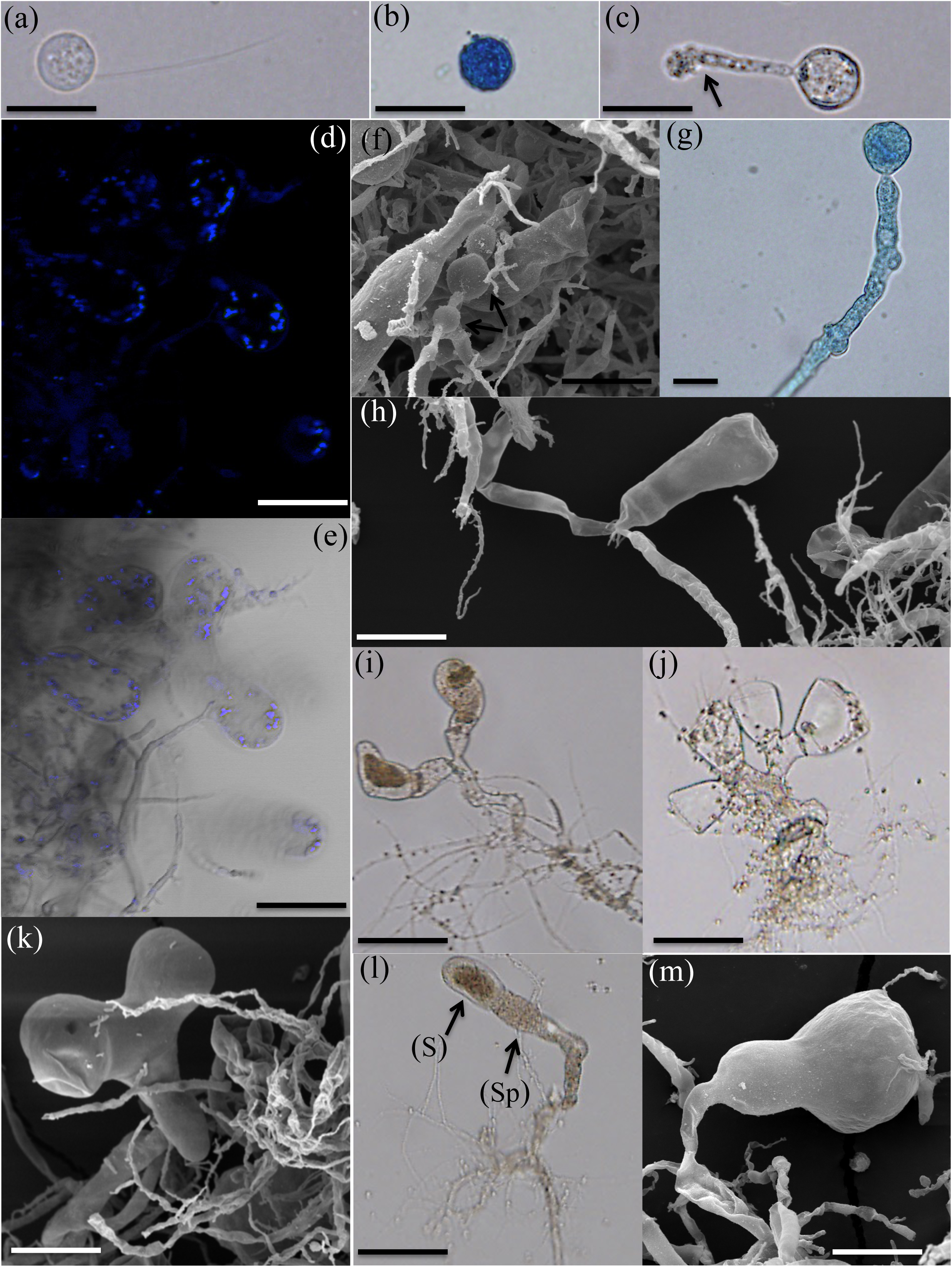
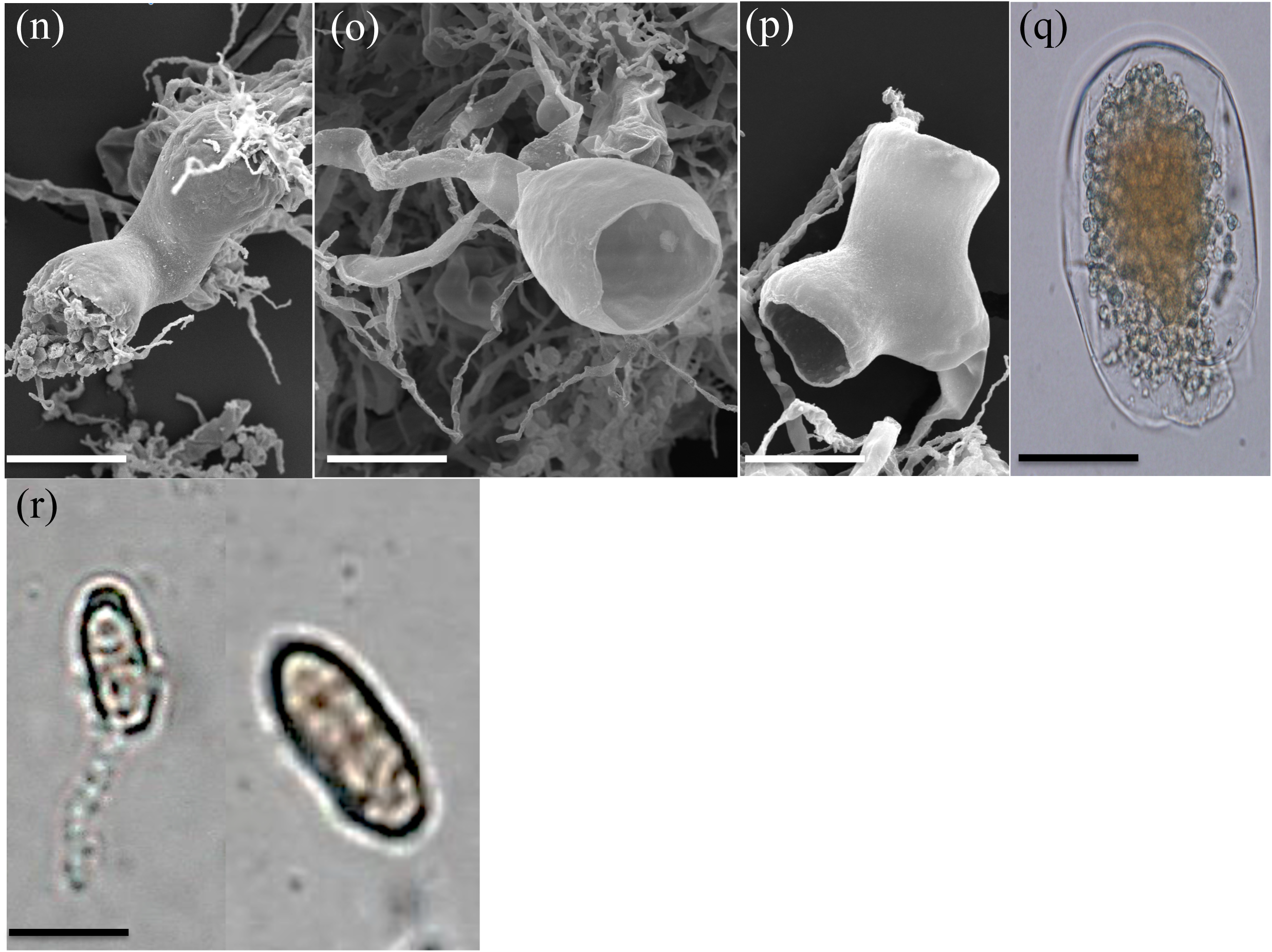
Microscopic features of *Khyollomyces ramosus* (Clade 6, Zebra-Horse) strain ZS-33 (a-q) and distinct resting stage structure from strain HoCal4.A2.2 (r). Light (a-e, g, i-j, l & q) and scanning electron (f, h, k, m &n-p) micrographs are shown. DAPI staining for nuclei visualizing using a fluorescence microscope equipped with a Brightline DAPI high-contrast filter set (d). Overlay image is shown in (e). (a) A uniflagellate zoospore. (b) Zoospore cyst after shedding of the flagellum. (c) Germinating zoospore cyst producing a germ tube (arrow). (d-e) Monocentric thalli, with nuclei occurring in sporangia, not in rhizoids or sporangiophores. (f) hyphal structures with intercalary swellings in wide hyphae (arrows). (g-h) Endogenous sporangial development: (g) Young subglobose sporangium with single rhizoidal system, (i) Mature ellipsoidal sporangium with two main rhizoidal systems. (i-m) Exogenous sporangial development: (i) Multisporangiate thallus with two sporangia, (j) Multisporangiate thallus with four sporangia, (k) Heart-shaped sporangium. (l) Ovoid sporangium (labeled S) on a wide flattened sporangiophore (labeled Sp), (m) Pyriform sporangium. (n) Zoospores are released through apical pore. (o-p) Empty sporangia with intact sporangial walls after zoospores discharge. (q) Mature sporangia detached from hyphae or sporangiophores. (r) Resting stages from strain HoCal4.A2.2. Bar: a-c & f-g =20 μm, d-e, k &m-q =50 μm and h-j &l =100 μm.

#### Clade 6 (Zebra-Horse) strains

Strain ZS-33 produced spherical zoospores, with an average diameter of 10.8 ± 3 μm (average ± standard deviation of 54 zoospores, range 6–17 μm). All zoospores were uniflagellate, with an average flagellum length of 26 ± 6.5 μm (average ± standard deviation of 54 zoospores, range 18–40 μm) (FIG. 8a). After shedding their flagella, zoospores started to encyst (FIG. 8b) and germinate producing germ tube (FIG. 8c). Germ tube branched and developed a highly branched anucleate rhizoidal system (FIG. 8d-e). Both narrow, 0.5–2.5 µm wide, and broad, 3–12.5 µm wide, hyphae were observed; intercalary swellings were frequently encountered in the broad hyphae (arrow in FIG. 8f).

Both endogenous and exogenous sporangia were observed. Endogenous sporangia varied in shape and size. Small endogenous sporangia were mainly subglobose (20–60 μm in diameter). (FIG. 8g). Large endogenous (80–160 μm L X 35–65 μm W) sporangia were mainly ellipsoidal (FIG. 8h). Exogenous sporangia size ranged between (80–270 μm L X 35–85 μm W) and displayed a wider range of morphologies, e.g. heart-shaped (FIG. 8k), ovoid (FIG. 8l) and pyriform (FIG. 8m). Sporangiophores ranged in length between 20–400 μm. Characteristically, strain ZS-33 displayed a multisporangiate thallus: the majority of sporangiophores were branched and bore two to four sporangia (FIG. 8i-j). Similar sporangial morphology has previously been observed in members of the genus *Piromyces* (e.g. *P. rhizinflatus*), and *Caecomyces* (e.g. *C. communis*) (Akin and others 1988; Akin and others 1989; Breton and others 1991). Unbranched sporangiophores with single sporangia were less frequently encountered (approximately 30% of observed sporangiophores n=50, Fig. 8k-m).

Zoospores were liberated through a wide apical pore at the top of the sporangia. The sporangial wall stayed intact after the discharge (FIG. 8n-p). Further, mature sporangia frequently detached from hyphae or sporangiophores, probably serving as an additional mean of fungal dispersal (FIG. 8q).

The type strain ZS-33 was obtained from Zebra fecal samples collected at the Oklahoma City Zoo. No noticeable differences were observed between ZS-33, and all other strains (n=15) obtained from Zebra fecal samples from the Oklahoma City Zoo. On the other hand, two distinct microscopic differences were identified in strains from domesticated Horses in Llanbadarn Fawr, Ceredigion County, Wales. First, multisporangiate thalli, copiously observed in ZS-33, were extremely rare in Welch Horse strains, and second, Distinct-resting stages (FIC. 8r) were often observed in Welch Horse strains, but never in Oklahoma City Zebra strains. Whether these differences are distinct characteristics of each group of strains, or induced by variations in media composition as well as growth and incubation procedures remain to be seen.

#### Clade 7 (Nilgiri Tahr) strains

Strain TDFKJa193 produced globose zoospores (FIG. 9a), with an average diameter of 3–4 μm. The majority of zoospores were mono-flagellate with 1, 2, or 3 flagella (FIG. 9a). Flagellum length ranged between 12–15 μm. Strain TDFKJa193 exhibited both endogenous and exogenous monocentric thallus development (FIG. 9b-e). Endogenous sporangia were terminal, varied in shape between globose, ovoid & obovoid, and ranged in size between 10–70 μm wide & 12–100 μm long (FIG. 9f-g). Some endogenous sporangia showed sub-sporangial swellings (FIG. 9f). Endogenous sporangia with one or two main rhizoidal systems (FIG. 9f) and with branched rhizoidal system (FIG. 9g) were also observed. Exogenous sporangia, on the other hand, were globose, ovoid & obovoid and were observed at the end of short sporangiophores (12–20 µm) (FIG. 9h-k). Some of the sporangiophores ended with sub-sporangial swellings with (FIG 9 i-j) or without (FIG. 9h&k) constricted neck of 1–8 μm width and 2–10 μm length. The presence of subsporangial swellings and short sporangiophores were previously reported for *Piromyces mae* (Ho and Barr 1995) and *Buwchfawromyces eastonii* (Callaghan and others 2015). Mature exogenous sporangia often showed the formation of a septum at their base (FIG. 9k) similar to *Neocallimastix frontalis* (Ho and Barr 1995). Zoospores liberation happened after irregular dissolution of the sporangial wall (FIG. 9l).

**FIG. 9.**
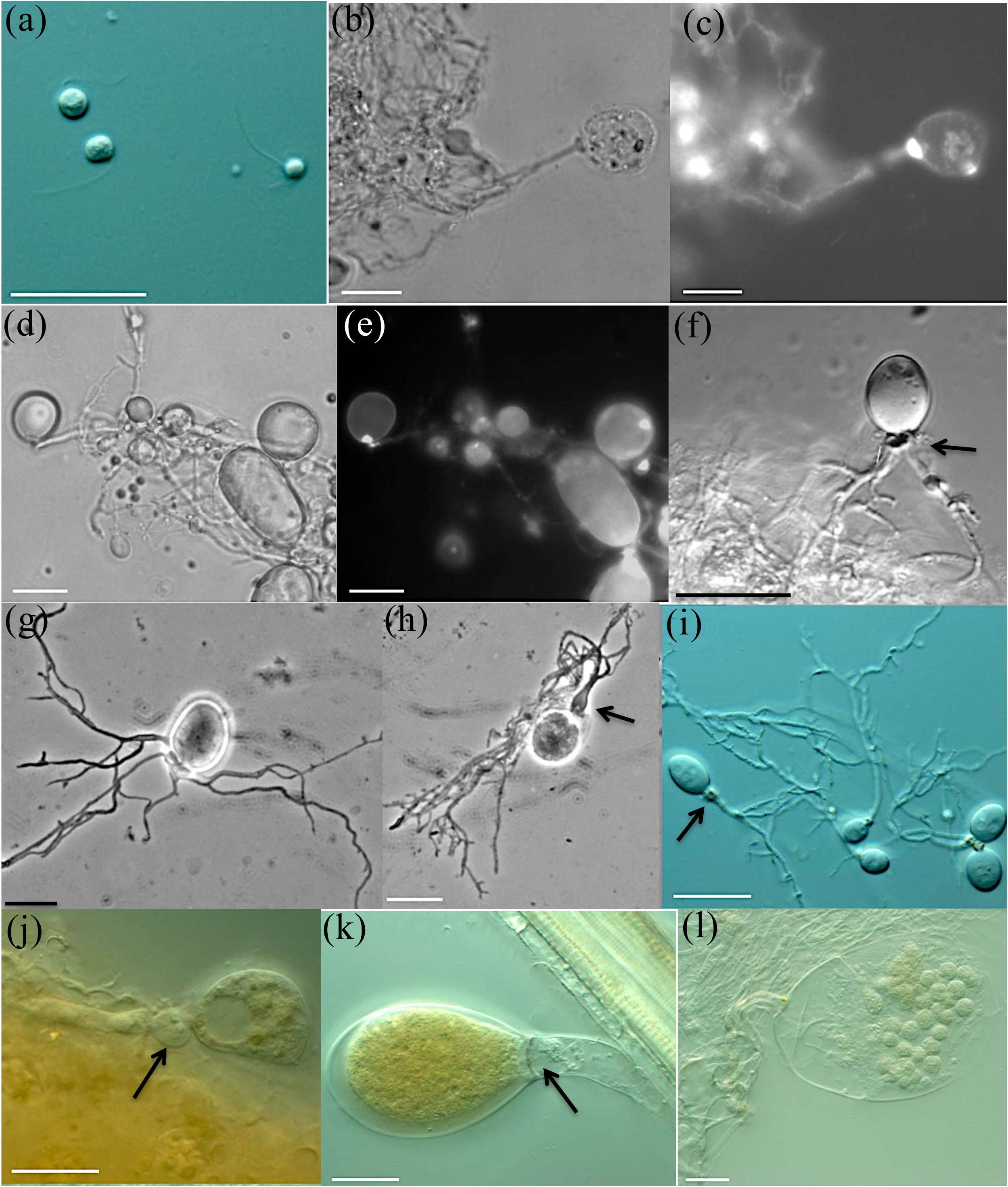
Microscopic features of *Tahromyces munnarensis* (Clade 7, Nilgiri Tahr) strain TDFKJa193. Differential interference contrast (a, f and i-l), phase contrast (b, d, g and h), fluorescence (c and e) micrographs. (a) A monoflagellate and triflagellate zoospores. (b-e) Monocentric thalli; nuclei were observed in sporangia, not in rhizoids or sporangiophores. (f-g) Endogenous sporangia. (f) Ovoid endogenous sporangium with two rhizoidal systems, note the sub-sporangial swelling (arrow). (g) Endogenous sporangium with branched rhizoids. (h-l) Exogenous sporangia: (h) Globose sporangium with short swollen sporangiophore (arrow). (i-j) Exogenous sporangia with sub-sporangial swellings and constricted necks (arrow). (k) Ovoid sporangium with septum at the sporangial base (arrow). (l) Zoospore release through dissolution of a wide apical pore. Scale bar = 20 µM.

### Phylogenetic analysis

Phylogenetic analysis using LSU (FIG. 10a) placed the isolated strains into seven monophyletic and bootstrap-supported lineages that were distinct from all currently described AGF genera. LSU sequence divergence estimates between various strains within a single clade ranged between 0–1%. The closest cultured representative to each of the seven clades is shown in TABLE 1. Within the LSU taxonomic framework, strains recovered from Axis Deer clustered within the *Orpinomyces-Neocallimastix-Pecoramyces-Feramyces* suprageneric clade, while strains recovered from Boer Goat and domesticated goat clustered within the *Oontomyces-Anaeromyces*-*Liebetanzomyces* supragenus clade. On the other hand, strains recovered from Nilgiri Tahr, as well as strains recovered from domesticated Goat and Sheep formed two distinct new clades associated with the genus *Buwchfawromyces*. In contrast, strains recovered from Zebra-Horse, and Mouflon-Boer Goat formed two distinct clades associated together but with no specific affiliation to any suprageneric group. The remaining group represented by strains isolated from White Tailed Deer displayed no specific affiliation to any currently characterized genera or supragenus groups within the Neocallimastigomycota.

**FIG. 10.**
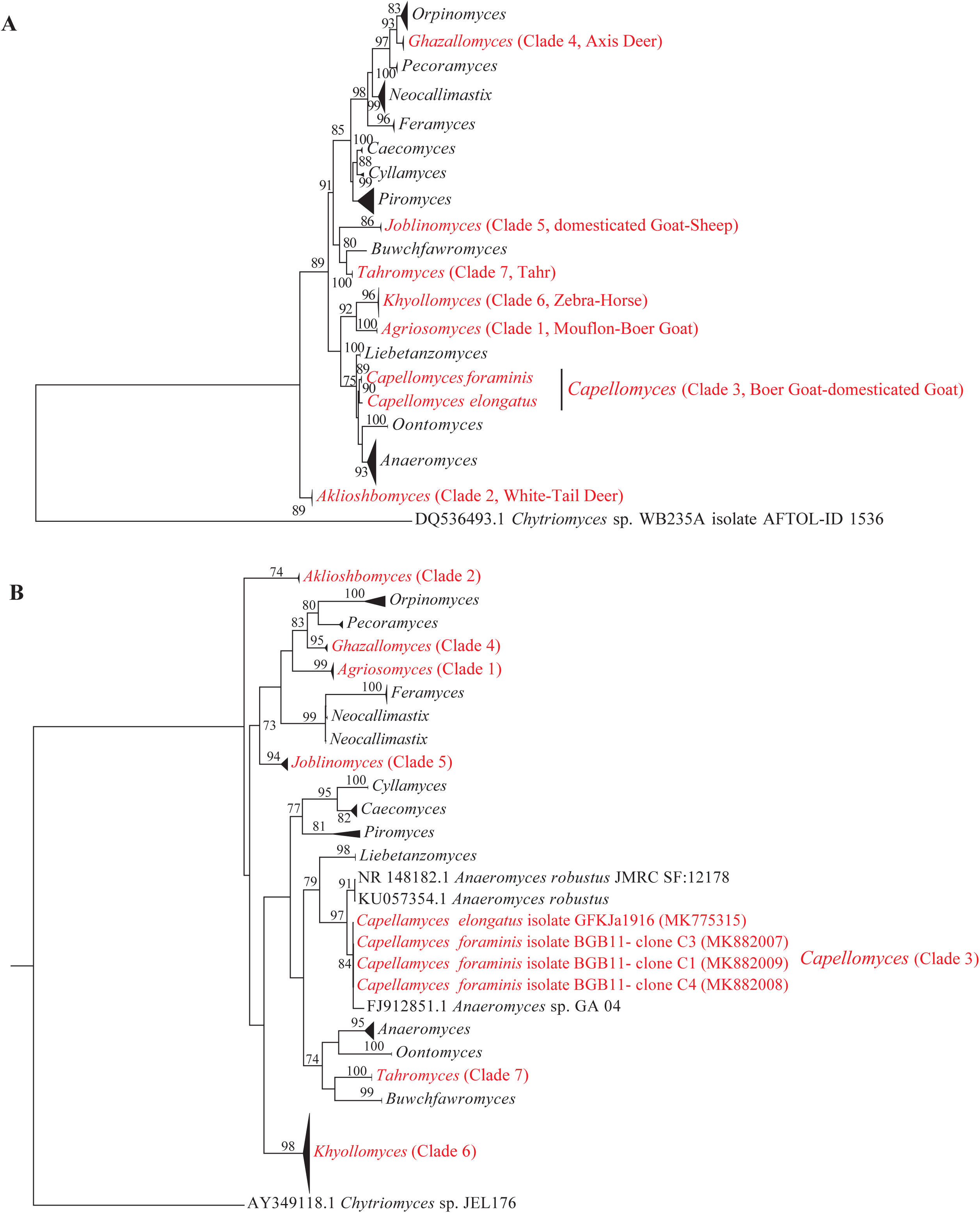
Phylogenetic affiliation of the 7 newly described genera to other AGF genera based on the sequences of the D1–D2 domains of nuc 28S rDNA gene (a), and partial ITS-1 sequences (b). Sequences were aligned in MAFFT (Nakamura and others 2018) and the alignment was used to construct phylogenetic trees in MEGA7 (Kumar and others 2016) using a maximum likelihood approach. Bootstrap values from 100 replicates are shown for nodes with more than 70% bootstrap support

To investigate ITS-1 sequence variability often reported within a single AGF strain, the ITS-1 region was amplified, cloned, and sequenced from all type strains. ITS-1 sequence divergence within type strains ranged between 0% for strain TDFKJa193 representative of the Nilgiri Tahr strains clade, and 0–8.4% (average 3.4%) for strain MS-2 representative of the Mouflon-Boer Goat strains clade. ITS-1-based analysis confirmed the monophyletic and distinct nature of all seven lineages, but yielded a different topology (FIG. 10b), as consistently observed in prior studies (Hanafy and others 2017; Wang and others 2017). Of special note, was the surprisingly high ITS-1 sequence similarity of the Boer Goat-domesticated Goat clade represented by strains BGB-11 and GFKJa1916 (FIG. 10B), to an *Anaeromyces* sp. isolate GA-04 (GenBank accession number FJ912851.1, unpublished) and to *Anaeromyces robustus* (GenBank accession number NR_148182.1 (Li and others 2016)). Average ITS-1 sequence divergence between various clones of the Boer Goat-domesticated Goat clade and *Anaeromyces* sp. GA-04 was 1.2%, and between various clones of the Boer Goat-domesticated Goat clade and *A. robustus* and was 4.2%. Unfortunately, the LSU sequence data from both *Anaeromyces* sp. GA-04 and *A. robustus* are not available for further comparison. This may be attributed to the inaccurate identification and reporting of these two cultures, an issue very well known in the anaerobic fungal taxonomy using ITS-1. We noted that in the paper describing *Anaeromyces robustus*, the type strain seems to deviate from the typical morphology of the genus *Anaeromyces*, e.g. multi-flagellate zoospores, lack of sausage shaped hyphae, and whale-tail like sporangia (see Figure 148 in (Li and others 2016)), casting doubts on the phylogenetic affiliation of *Anaeromyces robustus* to the genus *Anaeromyces*. Regardless, that stark morphological differences exist between Boer Goat-domesticated Goat strains and all other members of the genus *Anaeromyces*, e.g. monocentric thalli as opposed to polycentric thalli, absence of hyphal constrictions as opposed to sausage-shaped hyphae with multiple constrictions, and monoflagellate (2–4 flagella) versus uniflagellate zoospores (TABLE 1, FIG. 4-5). Such differences, in addition to the high ITS-1 sequence divergence values between the Boer Goat-domesticated Goat clade and other members of the genus *Anaeromyces* (7.1–13.1% to *A. mucronatus* and 8.5–18.6% to *A. contortus*), strongly support the distinction between these strains and the genus *Anaeromyces*.

### Ecological distribution

We queried the GenBank nr database to determine whether representatives of these seven novel clades were encountered in prior ITS-1-based culture-independent AGF diversity surveys. Multiple sequences with high (95.3-100%) sequence similarity to Zebra-Horse strain ZS-33 ITS-1 sequence were identified. These sequences were recovered from fecal samples obtained from multiple animals housed in the Oklahoma City Zoo (Liggenstoffer and others 2010), as well as in various locations (left and right dorsal colon, caecum and right ventral colon) within the digestive tract of horses (Mura and others 2019). This group has previously been assigned the alphanumeric designation (AL1) (Kittelmann and others 2012; Liggenstoffer and others 2010).

Surprisingly, ITS-1 sequences from the remaining six lineages (Axis Deer, White-Tailed Deer, Mouflon-Boer Goat, Boer Goat-domesticated Goat, domesticated Goat-domesticated Sheep, and Nilgiri Tahr) bore no close resemblance to all currently available ITS-1 sequence data (TABLE 1), with highest similarity being 83% in White-Tailed Deer to sequences from Bontebok, 84% in Mouflon-Boer Goat to sequences from Bontebok, 88% in domesticated goat-domesticated sheep to sequences from horse, 89% in Nilgiri Tahr strains to sequences from Okapi, 91% in Axis Deer to sequences from Llama, and 91-92% in Boer Goat-domesticated Goat strains to sequences from cow. As such, representatives of these novel lineages, the absolute majority of which (5/6) recovered from fecal samples of wild non-domesticated herbivores, do not appear to correspond to any of the alphanumerically designated uncultured groups previously identified in prior culture-independent efforts.

## TAXONOMY

***Agriosomyces*** Hanafy, Vikram B. Lanjekar, Prashant K. Dhakephalkar, T.M. Callaghan, Dagar, G.W. Griff, Elshahed, and N.H. Youssef, gen. nov.

MycoBank ID: MB830737

*Typification. Agriosomyces longus*. Hanafy, Vikram B. Lanjekar, Prashant K. Dhakephalkar, T.M. Callaghan, Dagar, G.W. Griff, Elshahed, and N.H. Youssef,

*Etymology: Agrioso*= derived from the Greek word for wild; *myces* = the Greek name for fungus. Obligate anaerobic fungus that produces small spherical monoflagellate zoospores with an extremely long flagellum (22 ± 3.8 μm). Zoospores germinate into monocentric thalli with filamentous anucleate rhizoidal systems. Both endogenous and exogenous globose sporangia are observed, which are very homogenous and display no pleomorphism. Rhizoids are swollen below the sporangial neck, which is tightly constricted. Zoospores are released through dissolution and rupturing of the sporangial wall. The clade is defined by the sequences MK882010-MK882013 (ITS-1) and MK881996 (D1-D2 28S rDNA).

***Agriosomyces longus*** Hanafy, Vikram B. Lanjekar, Prashant K. Dhakephalkar, T.M. Callaghan, Dagar, G.W. Griff, Elshahed, and N.H. Youssef, sp. nov.

MycoBank ID: MB830738.

*Typification*: The holotype is FIG. 2g in this manuscript, derived from the following: U.S.A. TEXAS: Val verde county, 29.369′ N and 100.829′ W ∼300 m ASL, 3d old culture of isolate MS-2, originally isolated from freshly deposited feces of male Mouflon Sheep (*Ovis orientalis*), April 2018, *Radwa Hanafy.* Ex-type strain: MS2, GenBank: MK881996 (D1-D2 28S rDNA).

*Etymology:* The species epithet “*longus”* refers to the extremely long flagellum observed in zoospores of strain MS-2 (FIG. 2a).

Obligate anaerobic fungus that produces small globose monoflagellate zoospores with an average diameter of 4 ± 1.1 μm. Zoospores are mainly mono-flagellate with a flagellum length of 22 ± 3.8 μm, approximately 5-6 times longer than the zoospore body. Bi-flagellate zoospores are rarely encountered. Zoospores germinate into monocentric thalli with filamentous anucleate rhizoidal systems. Both endogenous and exogenous sporangia are observed, which display no pleomorphism and both show globose morphology. In endogenous sporangia, the rhizoids are swollen below the sporangial neck, which is tightly constricted. Exogenous sporangia develop at the end of swollen sporangiophores, and the sporangial neck is constricted with a narrow neck port. Zoospores are released through dissolution and rupturing of the sporangial wall. Produces small brown spherical colonies on agar, and a thin biofilm-like growth in liquid media. The clade is defined by the sequences MK882010-MK882013 (ITS-1) and MK881996 (D1-D2 28S rDNA).

*Additional specimens examined:* Radwa Hanafy strain MS-4 (GenBank accession number of D1-D2 28S rDNA amplicon MK881997) isolated from the same freshly deposited feces of male Mouflon Sheep (*Ovis orientalis*) from which the type strain originated, April 2018, and strain BGS-13 (GenBank accession number of D1-D2 28S rDNA amplicon MK881995) isolated from freshly deposited feces of female wild Boer Goat *(Capra aegagrus)*, April, 2018.

***Aklioshbomyces*** Hanafy, Vikram B. Lanjekar, Prashant K. Dhakephalkar, T.M. Callaghan, Dagar, G.W. Griff, Elshahed, and N.H. Youssef, gen. nov.

MycoBank ID: MB830735

*Typification: Aklioshbomyces papillarum* Hanafy, Vikram B. Lanjekar, Prashant K. Dhakephalkar, T.M. Callaghan, Dagar, G.W. Griff, Elshahed, and N.H. Youssef.

*Etymology: Aklioshb*= derived from the Arabic word for grass-eaters (herbivores); *myces* = the Greek name for fungus.

Obligate anaerobic fungus that produces globose monoflagellate zoospores. Zoospores germinate into monocentric thalli with filamentous anucleate rhizoids. Exhibits both endogenous and exogenous monocentric thallus development. Exogenous sporangia develop at the end of unbranched sporangiophores of varying length. No morphological differences are observed between endogenous and exogenous sporangia, with ovoid, globose, and obpyriform sporangial shapes noted. The clade is defined by the sequences MK882038-MK882042 (ITS-1) and MK882001 (D1-D2 28S rDNA).

***Aklioshbomyces papillarum*** Hanafy, Vikram B. Lanjekar, Prashant K. Dhakephalkar, T.M. Callaghan, Dagar, G.W. Griff, Elshahed, and N.H. Youssef, sp. nov.

MycoBank ID: MB830736

*Typification*: The holotype is FIG. 3m in this manuscript, derived from the following: U.S.A.

OKLAHOMA: Payne county, 36.145′ N and 97.007′ W ∼300 m ASL, 3d old culture of isolate WT-2, originally isolated from freshly deposited feces of female White-Tailed Deer (*Odocoileus virginianus*), October 2017, *Radwa Hanafy.* Ex-type strain: WT-2, GenBank: MK882001 (D1-D2 28S rDNA).

*Etymology:* The species epithet “*papillarum”* refers to the papillae observed on the majority of strain WT-2 sporangia (FIG. 3m-q).

Obligate anaerobic fungus that produces globose monoflagellate zoospores with an average diameter of 7.4 ± 2.4 μm. The majority of zoospores are monoflagellate, with zoospores with two to three flagella less frequently observed. Fungal thalli are consistently monocentric with filamentous anucleate rhizoids. Germination of zoospores produces two types of monocentric thalli, endogenous and exogenous. Endogenous sporangia with single and two adjacent rhizoidal systems are observed. Pseudo-intercalary endogenous sporangia are occasionally observed. Sporangiophores carrying exogenous sporangia exhibit varying length from a few microns to 230 μm. Endogenous and exogenous sporangia are ovoid, globose, obpyriform, and ellipsoidal. Sporangia are mostly papillated with one or two papillae. Produces beige circular colonies with a brown central core of dense sporangial structures and an outer ring of light gray hyphal growth on agar, and heavy growth of thick biofilms that firmly attached to the tube’s glass surface in liquid media. The clade is defined by the sequences MK882038-MK882042 (ITS-1) and MK882001 (D1-D2 28S rDNA).

*Additional specimens examined:* Radwa Hanafy strains WT-1 (MK882000), WT-3 (MK881998), WT-4 (MK881999), WT-41(MK882002), WTS-51 (MK882006), WTS-52 (MK882003), WTS-53 (MK882004), and WTS-54 (MK882005), (GenBank accession number of D1-D2 28S rDNA amplicon in parenthesis) isolated from the same freshly deposited feces of female White-Tailed Deer from which the type strain originated, October 2017.

***Capellomyces*** Hanafy, Vikram B. Lanjekar, Prashant K. Dhakephalkar, T.M. Callaghan, Dagar, G.W. Griff, Elshahed, and N.H. Youssef, gen. nov.

MycoBank ID: MB830739.

*Typification. Capellomyces foraminis*. Hanafy, Vikram B. Lanjekar, Prashant K. Dhakephalkar, T.M. Callaghan, Dagar, G.W. Griff, Elshahed, and N.H. Youssef.

*Etymology: Capella*= derived from the Latin word for Goat; *myces* = the Greek name for fungus. Obligate anaerobic fungus that produces monoflagellate (1–3 flagella) zoospores. Zoospores germinate into monocentric thalli with filamentous anucleate rhizoidal systems. Both endogenous and exogenous sporangia are observed, with varying shapes and sizes. The clade is defined by the sequences MK882007-MK882009 (ITS-1), and MK881975 (D1-D2 28S rDNA). ***Capellomyces foraminis*** Hanafy, Vikram B. Lanjekar, Prashant K. Dhakephalkar, T.M. Callaghan, Dagar, G.W. Griff, Elshahed, and N.H. Youssef, sp. nov.

MycoBank ID: MB830740

*Typification*: The holotype is FIG. 4n in this manuscript, derived from the following: U.S.A.

U.S.A. TEXAS: Val verde county, 29.369′ N and 100.829′ W ∼300 m ASL, 3d old culture of isolate BGB-11, originally isolated from freshly deposited feces of a female Boer Goat (*Capra aegagrus*) April 2018, *Radwa Hanafy.* Ex-type strain: BGB-11, GenBank: MK881975 (D1-D2 28S rDNA).

*Etymology:* The species epithet “*foraminis”* refers to the wide apical pore at the top of the sporangia through which zoospores are discharged.

Obligate anaerobic fungus that produces spherical monoflagellate zoospores. Zoospores start to encyst after shedding their flagella. Zoospore cyst germinates, producing germ tube that subsequently branches into monocentric thalli with filamentous anucleate rhizoidal systems. Endogenous and exogenous sporangia are produced. Endogenous sporangia are ellipsoidal or ovoid. Exogenous sporangia are formed at the end of un-branched sporangiophores (20–150 μm). Some of the sporangiophores exhibit sub-sporangial swellings. Exogenous sporangia are ovoid, ellipsoidal with a single constriction, and globose. Zoospores are liberated through a wide apical pore at the top of the sporangia followed by sporangial wall collapse. Colonies are small (0.1–0.5 mm diameter) circular and brown, with dark center of sporangia structures on agar. Produces thin fungal biofilm in liquid media. The clade is defined by the sequences MK882007-MK882009 (ITS-1), and MK881975 (D1-D2 28S rDNA).

*Additional species examined.* Radwa Hanafy, strains BGB-2 (MK881974), BGC-12 (MK881976), BGS-11 (MK881977), and BGS-12 (MK881978) isolated from the same freshly deposited feces of a female Boer Goat (*Capra aegagrus*), April 2018, from which the type strain originated, April 2018 (D1-D2 28S rDNA amplicon GenBank accession number in parenthesis).

***Capellomyces elongatus*.** Hanafy, Vikram B. Lanjekar, Prashant K. Dhakephalkar, T.M. Callaghan, Dagar, G.W. Griff, Elshahed, and N.H. Youssef.

MycoBank: MB830869.

*Typification*: The holotype is FIG. 5k in this manuscript, derived from the following: India, KERALA, town of Munnar, 10.219′ N and 77.106′ E ∼2100 m ASL, 3d old culture of isolate GFKJa1916, originally isolated from freshly deposited feces of a domesticated but forest grazing Goat (*Capra aegagrus*), *Sumit S. Daggar.* Ex-type strain: GFKJa1916, GenBank: ITS-1 (MK775315), D1-D2 28S rDNA (MK775304).

*Etymology:* The species epithet “*elongatus”* refers to the characteristic long sporangiophore of exogenous sporangia.

Obligate anaerobic fungus that produces globose monoflagellate (1, 2, or 3 flagella) zoospores. Zoospore cyst germinate both endogenously and exogenously to produce monocentric thalli with filamentous anucleate rhizoidal systems. Endogenous sporangia are cylindrical, elongate, globose, sub-globose, ellipsoid & obovoid with sizes ranging between 8–60 μm wide & 10–140 μm long. Exogenous sporangia are formed at the end of developed at the end of long thick sporangiophores (up to 300 µm). Multisporangiate thalli are commonly observed with two sporangia of either the same or different shapes. Colonies are compact of 2–3 mm size, cottony and off-white in color with a compact and fluffy center made up of thick sporangia type structures, and surrounded by radiating rhizoids. Produces numerous fungal thalli that attach to the glass bottles on initial days of growth, which later develop into thin mat-like structures in liquid media. The clade is defined by the sequence MK775304 (D1-D2 28S rDNA).

*Additional species examined.* None

***Ghazallomyces*** Hanafy, Vikram B. Lanjekar, Prashant K. Dhakephalkar, T.M. Callaghan, Dagar, G.W. Griff, Elshahed, and N.H. Youssef, gen. nov.

MycoBank ID: MB830733

*Typification: Ghazallomyces constrictus* Hanafy, Vikram B. Lanjekar, Prashant K. Dhakephalkar, T.M. Callaghan, Dagar, G.W. Griff, Elshahed, and N.H. Youssef.

*Etymology: Ghazallo* = derived from the Arabic word for Deer (Ghazalla); *myces* = the Greek name for fungus.

Obligate anaerobic fungus that produces polyflagellate zoospores. Zoospores germinate into monocentric thalli with highly branched anucleate rhizoidal systems. Exhibits both endogenous and exogenous monocentric thallus development. Sporangia produced from endogenous and exogenous thalli development are pleomorphic, exhibiting a wide range of sporangial shapes.

During exogenous thallus development, zoospore cysts germinate from both ends, with rhizoids developing from one side and sporangiophore developing from the opposite side. The empty zoospore cyst remains as a persistent swollen structure at the base of unbranched sporangiophore that exhibit wide variations in lengths. Zoospores are released through an apical pore followed by collapse of the sporangial wall. The clade is defined by the sequences MK882043 (ITS-1) and MK881971 (D1-D2 28S rDNA).

***Ghazallomyces constrictus*** Hanafy, Vikram B. Lanjekar, Prashant K. Dhakephalkar, T.M. Callaghan, Dagar, G.W. Griff, Elshahed, and N.H. Youssef, sp. nov.

MycoBank ID: MB830734

*Typification*: The holotype is FIG. 6h in this manuscript, derived from the following: U.S.A. TEXAS: Sutton county, 30.591′ N and 100.138′ W ∼300 m ASL, 3d old culture of isolate Axs-31, originally isolated from freshly deposited feces content of female Axis Deer (*Axis axis)*, Apr. 2018, *Radwa Hanafy.* Ex-type strain: Axs-31, GenBank: MK881971 (D1-D2 28S rDNA).

*Etymology:* The species epithet “*constrictus”* refers to the observed constricted necks (point between sporangia and rhizoids) in the species endogenous sporangia (FIG. 6d-g).

Obligate anaerobic fungus that produces globose polyflagellate zoospores with 7–14 flagella. Zoospores germinate into monocentric thalli with highly branched anucleate rhizoidal systems. Exhibits both endogenous and exogenous monocentric thallus development. Endogenous sporangia produced from zoospore cyst enlargement develop into different shapes including globose, tubular, clavate, and ellipsoidal. Endogenous sporangia display tightly constricted necks (point between sporangia and rhizoids) with narrow ports. Exogenous sporangia develop at the end of unbranched sporangiophores of varied lengths. Both short (6–20μm) and long (up to 200μm) sporangiophores are observed. The exogenous sporangia display ellipsoidal, ovoid, globose, constricted ellipsoidal, pyriform, bowling pin-like, and rhomboidal shapes. Sporangial necks are constricted with narrow port. A fine septum develops at the base of the sporangium at maturity. Zoospores are released through an apical pore followed by collapse of the sporangial wall. Produces small circular white colonies (1–4 mm diameter) with a brown central core of dense sporangial structures on agar, and a thick fungal biofilm growth in liquid media. The clade is defined by the sequences MK882043 (ITS-1) and MK881971 (D1-D2 28S rDNA).

*Additional specimens examined:* Radwa Hanafy strains ADC-2 (MK881965), ADS-14 (MK881966), AXS-33 (MK881967), AXS-34 (MK1881968), ADS-12 (MK881969), AXS-32 (MK881970), ADS-11 (MK881972), and ADS-21 (MK881973) (GenBank accession number of D1-D2 28S rDNA amplicon in parenthesis) isolated from the same freshly deposited feces of female Axis Deer (*Axis axis)* from which the type strain originated, April 2018.

***Joblinomyces*** Hanafy, Vikram B. Lanjekar, Prashant K. Dhakephalkar, T.M. Callaghan, Dagar, G.W. Griff, Elshahed, and N.H. Youssef, gen. nov.

MycoBank ID: MB830867

*Typification. Joblinomyces apicalis*. Hanafy, Vikram B. Lanjekar, Prashant K. Dhakephalkar, T.M. Callaghan, Dagar, G.W. Griff, Elshahed, and N.H. Youssef.

*Etymology: Joblino*= honoring Keith N. Joblin for his contributions to the field of anaerobic fungi; *myces* = the Greek name for fungus.

Obligate anaerobic fungus that produces globose monoflagellate zoospores. Both endogenous and exogenous sporangia are observed with varying shapes and sizes. Sporangiophores of exogenous sporangia vary in length. Exogenous sporangia have short and frequently swollen sporangiophores. Zoospores discharge occurs through gradual dissolution of a wide apical portion of sporangial wall, resulting in formation of an empty cup-shaped sporangium. The clade is defined by the sequences MK910278 (ITS-1) and MK910268 (D1-D2 28S rDNA).

***Joblinomyces apicalis*.** Hanafy, Vikram B. Lanjekar, Prashant K. Dhakephalkar, T.M. Callaghan, Dagar, G.W. Griff, Elshahed, and N.H. Youssef.

MycoBank ID: MB830868

*Typification*: The holotype is FIG. 7c in this manuscript, derived from the following: India, HARYANA, city of Sonipat, 28.988′ N and 76.941′ E ∼220 m ASL, 3d old culture of isolate GFH683, originally isolated from freshly deposited feces of a domesticated goat (*Capra aegagrus hircus*), *Sumit Dagar.* Ex-type strain: GFH683, GenBank: MK910268 (D1-D2 28S rRNA).

*Etymology:* The species epithet “*apicalis”* refers to the zoospore discharge through the dissolution of a wide apical portion of the sporangial wall.

Obligate anaerobic fungus that produces globose monoflagellate zoospores with 1, or 2 flagella. Zoospores germinate to produce both endogenous and exogenous monocentric thalli.

Endogenous sporangia vary in shape between globose, sub-globose, ovoid, and obovoid with sizes ranging between 8–40 μm wide & 10–40 μm long. Exogenous sporangia are terminal and vary in shape between globose, ovoid, obovoid. Sporangiophores vary in length from 20–80 µm. Zoospores discharge occur through gradual dissolution of a wide apical portion of sporangial wall, resulting in formation of an empty cup-shaped sporangium. Produces 1–2 mm sized colonies with a dense dark central core of abundant sporangial growth, surrounded by long and thin radiating rhizoids. In liquid media, it produces numerous fungal thalli that attach to the glass bottles on initial days of growth, and later develop into thin mat-like structures. The clade is defined by the sequences MK910278-MK910282 (ITS-1) and MK910268-MK910272 (D1-D2 28S rDNA).

*Additional specimens examined:* Sumit Dagar strains GFH681 (MK910263-MK910267) and GFH682 (MK775330) (GenBank accession number of D1-D2 28S rDNA amplicon in parenthesis) isolated from the same freshly deposited domesticated goat feces from which the type strain originated, and SFH683 (MK775333) isolated from freshly deposited domesticated sheep feces in the city of Sonipat, Haryana, India.

***Khoyollomyces*** Hanafy, Vikram B. Lanjekar, Prashant K. Dhakephalkar, T.M. Callaghan, Dagar, G.W. Griff, Elshahed, and N.H. Youssef, gen. nov.

MycoBank ID: MB830741

*Typification. Khoyollomyces ramosus*. Hanafy, Vikram B. Lanjekar, Prashant K. Dhakephalkar, T.M. Callaghan, Dagar, G.W. Griff, Elshahed, and N.H. Youssef.

*Etymology: Khyollo*= derived from the Arabic word for horses; *myces* = the Greek name for fungus.

Obligate anaerobic fungus that produces spherical uniflagellate zoospores. Zoospores encyst and develop a highly branched anucleate rhizoidal system. Both endogenous and exogenous sporangia are observed. Small endogenous sporangia are subglobose and large endogenous sporangia are ellipsoidal. Exogenous sporangia displayed a wider range of shapes. The majority of sporangiophores are branched and bear two to four sporangia. Unbranched sporangiophores bearing a single sporangium are less frequently encountered. Zoospores are liberated through a wide apical pore at the top of the sporangia. Mainly found in the digestive tracts of equids. The clade is defined by the sequences MK882019 (ITS-1), and MK881981 (D1-D2 28S rDNA).

***Khoyollomyces ramosus*.** Hanafy, Vikram B. Lanjekar, Prashant K. Dhakephalkar, T.M. Callaghan, Dagar, G.W. Griff, Elshahed, and N.H. Youssef, sp. nov.

MycoBank ID: MB830742.

*Typification*: The holotype is FIG. 8j in this manuscript, derived from the following: U.S.A. OKLAHOMA: Oklahoma City, 35.524′ N and 97.472′ W ∼300 m ASL, 3d old culture of isolate ZS-33, originally isolated from freshly deposited feces of a Grevy’s Zebra *(Equus grevyi)*, May, 2018, *Radwa Hanafy.* Ex-type strain: ZS-33, GenBank: MK881981 (D1-D2 28S rDNA).

*Etymology:* The species epithet “*ramosus”* (Latin for branched) refers to the observed branched sporangiophores bearing two to four sporangia in *K. ramosus* type strain ZS-33 (FIG. 8i-k).

Obligate anaerobic fungus that produces spherical uniflagellate zoospores. Zoospores encyst and germinate producing germ tube that develops into a highly branched anucleate rhizoidal system. Both narrow, 0.5–2.5µm wide, and broad hyphae, 3–12.5µm wide, are produced; intercalary swellings are frequently encountered in the broad hyphae. Both endogenous and exogenous sporangia were observed. Endogenous sporangia vary in shape and size, with small endogenous sporangia mainly subglobose (20–60 μm in diameter) while large endogenous (80–160μm L X 35–65μm W) sporangia mainly ellipsoidal. Exogenous sporangia ranged in size between (80– 270μm L X 35–85μm W)) and display a wide range of morphologies, e.g. heart-shaped, ovoid, and pyriform. Displays a multisporangiate thallus, with the majority of sporangiophores being branched and bearing two to four sporangia. Unbranched sporangiophores with single sporangia are less frequently encountered (approximately 30% of observed sporangiophores). Zoospores are liberated through a wide apical pore at the top of the sporangia. The sporangia stay intact after the discharge. Mature sporangia frequently detach from hyphae or sporangiophores.

Produces small yellow to yellowish brown irregularly shaped colonies on agar. In liquid media, the fungal growth is loose and exhibited a sand-like appearance. The clade is defined by the sequences MK882019 (ITS-1), and MK881981 (D1-D2 28S rDNA).

*Additional specimens examined:* Radwa Hanafy strains ZC-31 (MK881979), ZC-32 (MK881980), ZC-33 (MK881981), ZC-41 (MK881982), ZC-42 (MK881983), ZC-43 (MK881984), ZC-51 (MK881985), ZC-53 (MK881986), ZS-21 (MK881987), ZS-22 (MK881988), ZS-31 (MK881989), ZS-32 (MK881990), ZS-41 (MK881992), ZS-42 (MK881993), and ZS-43 (MK881994) (GenBank accession number of D1-D2 28S rDNA amplicon in parenthesis) isolated from the same freshly deposited Zebra feces from which the type strain originated, May 2018. Tony Callaghan strains: HoCal4.A2, HoCal4.A2.2, HoCal4.A4 isolated from fresh horse feces (Llanbadarn, nr. Aberystwyth; 52.4156,-3.8878), August 2013. Two further cultures (Tmc003.6a, TMC3.6b) were isolated from a different horse at the same site, November 2013.

***Tahromyces*** Hanafy, Vikram B. Lanjekar, Prashant K. Dhakephalkar, T.M. Callaghan, Dagar, G.W. Griff, Elshahed, and N.H. Youssef, gen. nov.

MycoBank ID: MB830865

*Typification. Tahromyces munnarensis*. Hanafy, Vikram B. Lanjekar, Prashant K. Dhakephalkar, T.M. Callaghan, Dagar, G.W. Griff, Elshahed, and N.H. Youssef.

*Etymology: Tahro*= referring to the Nilgiri Tahr from which the species was isolated; *myces* = the Greek name for fungus.

Obligate anaerobic fungus that produces globose monoflagellate zoospores. Both endogenous and exogenous sporangia are observed with varying shapes and sizes. Endogenous sporangia with one or two main rhizoidal systems and with branched rhizoidal system are frequently observed. Exogenous sporangia have short and frequently swollen sporangiophores. Sporangial necks are frequently constricted. Septa often form at the base of mature exogenous sporangia. Zoospores liberation happens after irregular dissolution of the sporangial wall. The clade is defined by the sequences MK775321 (ITS-1), and MK775310 (D1-D2 28S rDNA).

***Tahromyces munnarensis*** Hanafy, Vikram B. Lanjekar, Prashant K. Dhakephalkar, T.M. Callaghan, Dagar, G.W. Griff, Elshahed, and N.H. Youssef.

MycoBank ID: MB830866

*Typification*: The holotype is FIG. 9h in this manuscript, derived from the following: India, KERALA, town of Munnar, 10.219′ N and 77.106′ E ∼2100 m ASL, 3d old culture of isolate TDFKJa193, originally isolated from freshly deposited feces of a Nilgiri Tahr *(Nilgiritragus hylocrius), Sumit Dagar.* Ex-type strain: TDFKJa193, GenBank: ITS-1 accession number (MK775321), D1-D2 28S rDNA (MK775310).

*Etymology:* The species epithet “*munnarensis”* refers to the town that the type species was isolated from.

Obligate anaerobic fungus that produces globose monoflagellate zoospores with 1, 2, or 3 flagella. Both endogenous and exogenous sporangia were observed. The sporangia vary in size between 12–100 μm in length and 10–70 μm in width, and display a wide range of morphologies like globose, ovoid, and obovoid. Sporangiophores are short (12–20 µm) with frequent subsporangial swellings. Sporangial necks (1–8 μm width and 2–10 μm length) are frequently constricted. Septa often form at the base of mature exogenous sporangia. Zoospores liberation happens after irregular dissolution of the sporangial wall. Produces colonies that are small (1 mm), white in color with a compact and fluffy center, surrounded by dotted circles of fungal thalli. In liquid media, it produces numerous fungal thalli attaching to the glass bottles on initial days of growth, and later developing into thin mat-like structures. The clade is defined by the sequence MK775310 (D1-D2 28S rDNA).

*Additional specimens examined:* Sumit Dagar strains TDFKJa1924 (MK775323), TDFKJa1926 (MK775322), and TDFKJa1927 (MK775324) (GenBank accession number of D1-D2 28S rDNA amplicon in parenthesis) isolated from the same Nilgirir Tahr feces from which the type strain originated.

## DISCUSSION

Here, we report on the isolation and characterization of multiple novel AGF strains from a concerted sampling effort of domesticated, zoo-housed, and wild animals from North America, Europe, and Asia. We propose seven new AGF genera to accommodate these novel strains, hence expanding the AGF genus-level diversity by more than 50% (From 11 to 18). All newly described taxa produced filamentous, monocentric thalli, similar to seven of the eleven currently described genera. Six of the seven novel genera described here produce mono/uniflagellate zoospores, similar to eight of the eleven currently described taxa. As such, filamentous taxa with moncentric thalli and monflagellate zoospore appear to be the most common thallus morphology and zoospore flagellation patterns in the Neocallimastigomycota predominant within currently described AGF genera. It is interesting to note that for decades, microscopic-based identification of AGF strains typically assigned isolates with such morphology to the genus *Piromyces* (Ho and others 1993). We note broad similarities between the microscopic features of *Aklioshbomyces papillarum and P. mae* (papillated sporangia), and *Joblinomyces* and *P. Minutus* (zoospores release through a wide apical portion of sporangial wall, resulting in formation of an empty cup-shaped sporangium). Unfortunately, the absnece of sequence data and extant cultures of these previously described “*Piromyces*” taxa prevents futher investigation into this issue (Ho and others 1993).

The current isolates were obtained in a multi-year effort to describe novel AGF strains from a wide range of animal hosts in the United States, India, Wales. The majority of novel taxa described here (5/7 genera, 6/8 species) originated from wild undomesticated animals (Axis Deer, White Tailed Deer, Mouflon, Boer Goat, and Nilgiri Tahr), underscoring their potential as novel, yet-untapped reservoir of AGF diversity. Such novelty, which has recently been postulated (Hanafy and others 2018) could be attributed to higher variability in the quality and quantity of ingested plant material, and the significant daily and seasonal fluctuations in feeding frequencies.

Culture-independent surveys utilizing ITS-1 as a phylogenetic marker (Kittelmann and others 2012; Liggenstoffer and others 2010), and subsequent meta-analysis (Kittelmann and others 2012; Kittelmann and others 2013) have identified multiple novel yet-uncultured AGF genus-level lineages. This study has been successful in isolating the first representatives of novel group AL1 from Zebra and Horse fecal samples. In a prior survey of AGF in zoo-housed animals (Liggenstoffer and others 2010), members of this lineage were encountered in approximately half of the animal hosts examined (18/35). AL1-affiliated sequences were more predominant in hindgut fermenters (7/9 hosts), and comprised a relatively high proportion of the AGF community in multiple hosts, e.g. 99.9% in three different Zebra individuals, 56.7 and 68.3% in two horses, and 29.6% in a Grant’s Gazelle). By comparison, they were only encountered in 11/26 foregut fermenters, where they constituted 0.01% to 38% of the AGF community in these animals. Further a recent seminal spatial analysis that analyzed AGF community in samples directly obtained from various locations along horses digestive tracts (Mura and others 2019) identified AL1 group as a prominent component of the AGF community in the right ventral (88%), and left dorsal (98%) colons in horses. As such, this novel genus appears to exhibit a preference for hindgut fermenters of the family Equidae. The reason for such preferences, and the general preference of some fungal taxa to specific hosts remains unclear (Callaghan and others 2015; Dagar and others 2015).

Surprisingly, comparative analysis of ITS-1 sequences indicated that the Axis Deer, White tailed Deer, Mouflon-Boer Goat, Boer Goat-domesticated Goat, Nilgiri Tahr, and domesticated Goat-sheep groups appear to be completely novel, and previously unencountered in prior culture-based or culture-independent studies. ITS-1 sequences from these four isolates did not display mismatches to common ITS-1 primers, did not have an atypical length that could hinder its amplification or detection via PCR, and were readily amplified from pure-cultures’ genomic DNA. As such, we posit that the lack of prior observation of these taxa is biologically relevant, and is indicative of their relatively specific host preference and/or predominance in wild, rather than domesticated herbivores. Indeed, although 30 animals were screened in the current study, three of these seven novel genera were isolated only from a single host (White tailed Deer, Axis Deer, and Nilgiri Tahr), while the other four were isolated from only two hosts (Mouflon and Boer Goat, Boer Goat and domesticated Goat, domesticated Goat and Sheep, and Zebra-domesticated Horse) (TABLE 1).

Collectively, the steady identification of novel taxa in culture-based and culture-independent surveys, as well as the sparse overlap between these studies strongly suggests that the scope of AGF diversity in nature is much broader than currently estimated (Kittelmann and others 2012; Paul and others 2018). Compared to the prokaryotic component of the rumen and herbivorous gut, the diversity of the rumen mycobiome remains woefully understudied. To provide a more thorough understanding of the AGF diversity in nature, concerted efforts that systematically assess the AGF diversity and community structure in various spatial (e.g. across various compartments of the herbivorous gut), temporal (e.g. across the lifespan of an animal), and geographic dimensions in a wide range of domesticated and wild herbivores is needed. Much remains to be understood regarding the diversity and community structure of AGF within various locations of the gastrointestinal tract of an animal host, interspecies stochastic differences between AGF communities in animal subjects, temporal age-related progression of AGF in animal hosts, and the response of the AGF community to various factors e.g. feeding patterns, antibiotic administration, animal disease, and co-housing arrangements and combinations thereof.

## ACKNOWLEDGMENTS

We thank Jim and Tammy Austin for providing fecal samples, Britny Johnson for technical assistance, Dr. Karthick Balasubramanian for his valuable help, Drs. Rebecca Snyder and Jennifer D’Agostino for providing fecal samples from the Oklahoma City Zoo, and to the Kerala Forest Department, Munnar for sampling permission (WL10-51713/2017). This work has been funded by the NSF-DEB Grant number 1557102 to NHY and MSE, Department of Biotechnology (DBT) project no. BT/PR15694/PBD/26/506/2015 to SSD.

